# Total Synthesis of Self-Assembling Semi-Synthetic Proteins Utilizing a Dendritic Solubility Tag

**DOI:** 10.64898/2026.07.24.740575

**Authors:** Kshitish C Hati, Britto S. Sandanaraj

## Abstract

The chemical synthesis of well-defined, self-assembling semi-synthetic proteins (SSPs) has attracted growing interest in recent years. Approaches such as micelle-assisted protein-labeling technology (MAPLabTech) and supramolecule-assisted protein-labeling technology (SAPLabTech) have been used to generate a wide range of SSPs. The central challenge in synthesizing SSPs is solubilizing a hydrophobic chemical probe in aqueous medium prior to bioconjugation. Both MAPLabTech and SAPLabTech rely on non-covalent interactions to solubilize hydrophobic probes and present certain limitations. The present study introduces a complementary chemical strategy in which a hydrophobic chemical probe is covalently tagged with a cleavable, water-soluble dendritic domain. This covalent tagging renders the probe fully water-soluble, enabling quantitative bioconjugation to yield monomeric semi-synthetic proteins. Subsequent, selective removal of the solubility tag converts the hydrophilic semi-synthetic proteins into facially amphiphilic, semi-synthetic proteins.

## Introduction

Self-assembling proteins (SAPs) have attracted considerable interest in recent years for their utility across a range of biomedical applications.^1–6^ They are generated primarily through biological^7–9^ or chemical methods.^10,11^ Biological methods have become powerful, robust, and mature technologies for the custom design of SAPs; however, this approach relies on only the twenty naturally occurring amino acids, which limits functional diversity.^7–9^ On the other hand, chemical technologies offer unlimited possibilities of building blocks to construct SAPs.^12,13^ Although extremely powerful approach, chemical methods, by contrast, remain comparatively immature. SAPs generated through chemical synthesis are often poorly defined and polydisperse, and have generally lacked rigorous structural and biophysical characterization.^14^

Our laboratory entered this field in 2014 with the goal of designing and synthesizing well-defined SAPs possessing rich structural and functional diversity. Toward that end, we developed micelle-assisted protein-labeling technology (MAPLabTech)^15^**(Figure 1a)**, which has since been used over the past decade to synthesize functional SAPs with an exceptional precision^16–22^. More recently, we introduced a complementary approach, supramolecule-assisted protein-labeling technology (SAPLabTech)^23^**(Figure 1b)**. Both technologies rely on non-covalent interactions to solubilize hydrophobic chemical probes, enabling efficient bioconjugation to generate SSPs.^15–23^

**Figure 1.**
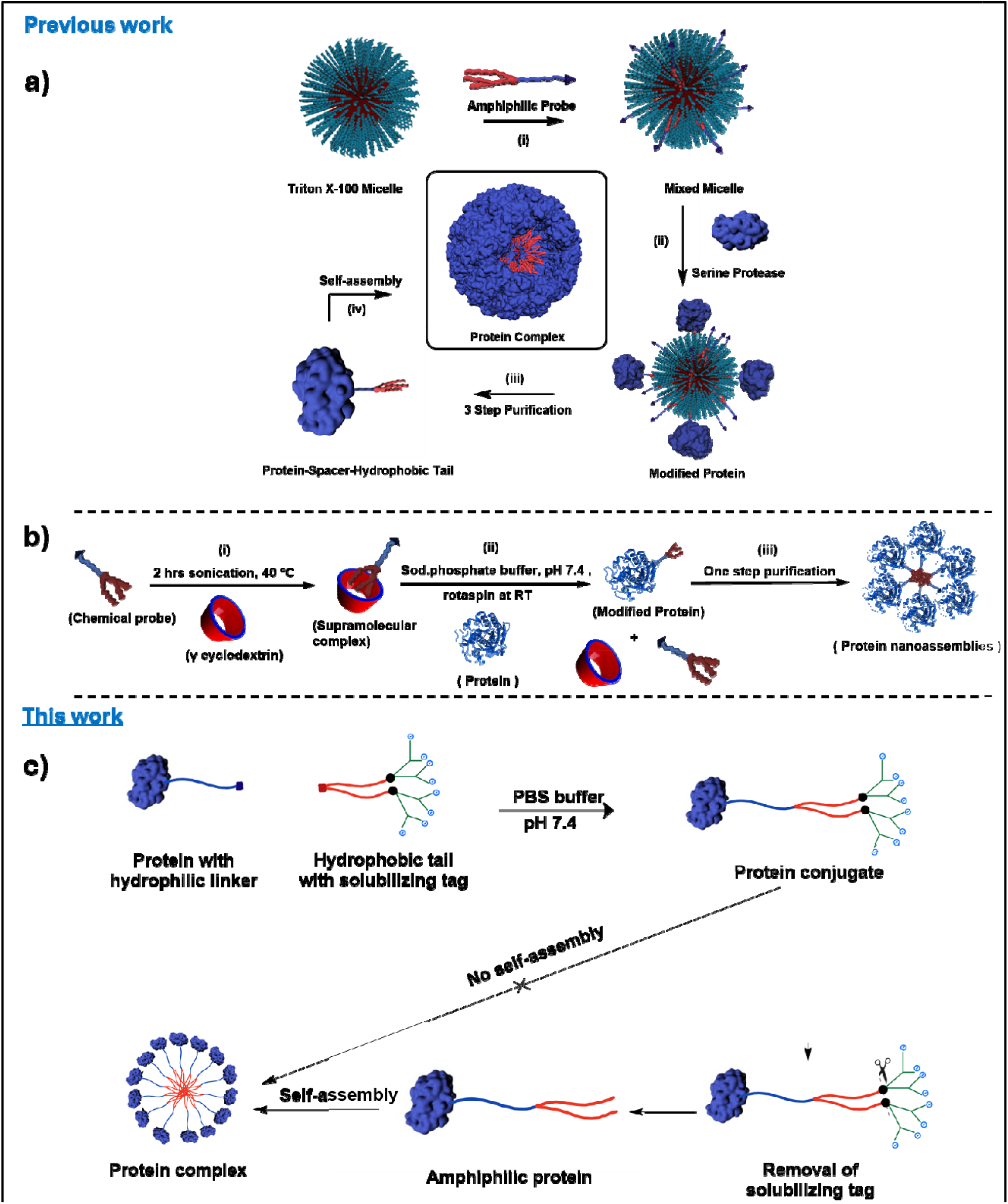
Cartoon representation of; a) Micelle assisted protein labelling technology (MAPLabTech), b) Supramolecule assisted protein labeling technology (SAPLabTech), c) Solubilizing tag-assisted protein labeling technology

The present study expands this chemical toolbox by introducing an orthogonal, complementary method, outlined conceptually in **Figure 1c**. In this approach, a hydrophobic chemical probe is rendered highly hydrophilic through attachment of a cleavable solubility tag. This modification enables quantitative bioconjugation to produce a well defined, monodisperse, hydrophilic protein bioconjugate. Selective cleavage of the solubility tag subsequently yields a facially amphiphilic, self-assembling semi-synthetic protein. Herein, we describe the total synthesis of self-assembling semi-synthetic proteins (SSPs).

## Results

### Molecular Design of Target Probe

The ideal probe should comprise five essential structural elements: (i) a protein-reactive head group for site-selective conjugation, (ii) a hydrophilic spacer to ensure aqueous compatibility and provide spatial separation from the protein surface, (iii) a hydrophobic group responsible for self-assembly, (iv) a stimuli-responsive linker enabling controlled activation or release, and (v) a solubilizing tag to solubilize the hydrophobic tail. The simultaneous incorporation of these chemically distinct functionalities into a single molecular scaffold presents a considerable synthetic challenge. Their differing reactivities necessitate multiple protection and deprotection strategies, resulting in lengthy synthetic routes, diminished overall yields, and limited flexibility for structural diversification. To streamline the synthetic workflow and maximize modularity, we developed a convergent approach wherein the native protein was first functionalized with a cyclooctyne-terminated linker. In parallel, the hydrophobic tail was designed to incorporate an azide terminus and was functionalized with a peripheral carboxylic acid-terminated dendron through a redox-cleavable disulfide linker. This design ensures that the hydrophobic domain remains highly soluble in aqueous buffer during all handling stages.

### Synthesis of Activated Dendritic Ester

The synthesis of the dendritic intermediate was carried out through a multistep sequence involving selective functionalization and activation reactions. Initially, 3,5-dihydroxybenzyl alcohol was selectively O-alkylated with *tert*-butyl bromoacetate in the presence of potassium carbonate (K_2_CO_3_) as a base, affording the corresponding *tert*-butyl ester intermediate **1a** in good yield. The hydroxymethyl group of **1a** was then converted into the corresponding benzyl bromide **1b** through bromination using tetrabromomethane (CBr_4_) and triphenylphosphine (PPh_3_) in dichloromethane (DCM) under mild conditions. The resulting benzyl bromide **1b** was reacted with 3,5-dihydroxybenzyl alcohol to furnish the dendritic alcohol **1c**. Finally, the terminal hydroxyl group of **1c** was activated by reaction with *p*-nitrophenyl chloroformate (*p*-NPC), leading to the formation of the corresponding reactive dendritic carbonate ester **1d**. The p-nitrophenyl carbonate serves as a chemoselective reactive handle for carbamate bond formation with amine-containing substrates, making **1d** a versatile precursor for the preparation of solubilizing tags **(Scheme 1)**.

### Synthesis of Azide-terminated Disulfide-functionalized Hydrophobic Chemical Probe

The azide-terminated, disulfide-functionalized hydrophobic intermediate **2h** was synthesized through a convergent multistep strategy involving selective functional group manipulations, orthogonal protection, and efficient amide bond formation. The synthesis commenced with the Fischer esterification of 5-hydroxyisophthalic acid in methanol under catalytic sulfuric acid, affording the corresponding dimethyl ester **2a** in excellent yield. Controlled partial reduction of **2a** with lithium aluminium hydride (LiAlH_4_) in anhydrous tetrahydrofuran (THF) enabled the chemoselective reduction of a single methyl ester, furnishing the hydroxymethyl monoester **2b** while preserving the second ester functionality for subsequent derivatization. The phenolic hydroxyl group of **2b** was then selectively O-alkylated with 1-bromooctadecane in the presence of potassium carbonate (K_2_CO_3_), providing the octadecyl ether derivative **2c** in good yield. The benzylic hydroxymethyl group of **2c** was subsequently activated by mesylation using methanesulfonyl chloride under standard conditions. The resulting mesylate intermediate **2d** was obtained in sufficient purity and was carried forward directly without chromatographic purification. Subsequent nucleophilic substitution with sodium azide in DMF proceeded smoothly to furnish the azide-terminated monoester **2e** in high yield, thereby installing a bioorthogonal functional handle suitable for subsequent copper(I)-catalyzed azide–alkyne cycloaddition (CuAAC) ligation. Hydrolysis of the methyl ester under basic conditions using sodium hydroxide in methanol afforded the corresponding carboxylic acid **2f** quantitatively, which was employed directly in the next step without further purification. In parallel, selective mono-*N*-Boc protection of cystamine dihydrochloride was accomplished to provide the unsymmetrically protected disulfide linker **2g** in excellent yield while retaining one free primary amine for chemoselective coupling. Finally, HATU-mediated coupling between obtained carboxylic acid derivative **2f** and mono-*N*-Boc-protected cystamine **2g** efficiently afforded the target intermediate **2h** in good yield. The convergent synthetic design integrates three key structural motifs-a hydrophobic C18 alkyl chain, a bio-orthogonal azide functionality, and a reduction-responsive disulfide linker within a single molecular framework, thereby furnishing a versatile precursor for the modular construction of stimuli-responsive dendritic probes. **(Scheme 2)**

### Synthesis of an Azide-terminated, Disulfide-functionalized Hydrophilic Chemical Probe

Treatment of azide intermediate **2h** with trifluoroacetic acid (TFA) in dichloromethane (DCM) quantitatively removed the *N*-Boc protecting group, affording the corresponding free amine. Owing to the inherent instability of the deprotected amine **3a**, the crude product was used directly in the subsequent coupling reaction without further purification. The resulting amine was reacted with the activated dendritic carbonate ester **1d** in DCM to furnish the dendritic ester-based azide **3b**, incorporating the disulfide linker in good yield. The dendritic intermediate **3b** was subsequently conjugated to the previously reported tetraethylene glycol (TEG)-based tosylated mono-propargylated building block **3c**^15^ through a copper(I)-catalyzed azide–alkyne cycloaddition (CuAAC) in THF/H_2_O. The click reaction proceeded efficiently and with excellent chemoselectivity, affording the triazole-linked intermediate **3d** in excellent yield. The tosylate intermediate **3d** was subsequently converted to the corresponding azide **3e** by treatment with sodium azide in DMF under standard azidation conditions. This transformation proceeded in high yield and introduced an additional orthogonal azide functionality for subsequent derivatization. Finally, the tert-butyl protected carboxylate moiety of **3e** was deprotected using TFA in DCM, and the resulting product was isolated by precipitation with diethyl ether to furnish the target terminal dendritic acid based amphiphilic molecule **3f** in high purity and excellent overall yield. This modular synthetic sequence, comprising sequential deprotection, carbamate formation, CuAAC ligation, nucleophilic azidation, and final ester deprotection, highlights the versatility of the approach for the efficient construction of multifunctional amphiphilic dendritic architectures containing both disulfide and azide functionalities. **(Scheme 3)**

### Bio-conjugation of COT-terminated Linker with Human Serum Albumin (HSA)

To evaluate the feasibility of the bioconjugation strategy, human serum albumin (HSA) was selected as the initial model protein. HSA is one of the most widely employed protein scaffolds in bioconjugation studies owing to its high aqueous solubility, exceptional stability, and well-characterized three-dimensional structure.^24,25^ Importantly, HSA contains a single solvent-accessible free cysteine residue (Cys34), whereas the remaining cysteine residues are engaged in intramolecular disulfide bonds.^26,27^ This unique reactivity enables highly site-selective thiol modification under mild conditions, making HSA an ideal model for assessing the efficiency and chemoselectivity of the developed conjugation methodology.^28^ Site-selective modification of native human serum albumin (HSA) was achieved by targeting the solvent-accessible Cys34 residue with a cyclooctyne (COT)-maleimide probe (2 eq.) under mild aqueous conditions, affording the COT-functionalized HSA conjugate for subsequent bioorthogonal conjugation studies. Reaction progress was monitored by MALDI-TOF mass spectrometry, which showed a shift from the native protein peak at m/z 66,654 Da to a single product peak at m/z 67,233 Da, indicating efficient and homogeneous modification **(Figure 2)**. This mass shift of approximately 600 Da quantitatively matches the theoretical monoadduct stoichiometry, confirming complete and homogeneous conversion to the HSA-COT conjugate without detectable multi-labeling or unreacted precursor.

**Figure 2.**
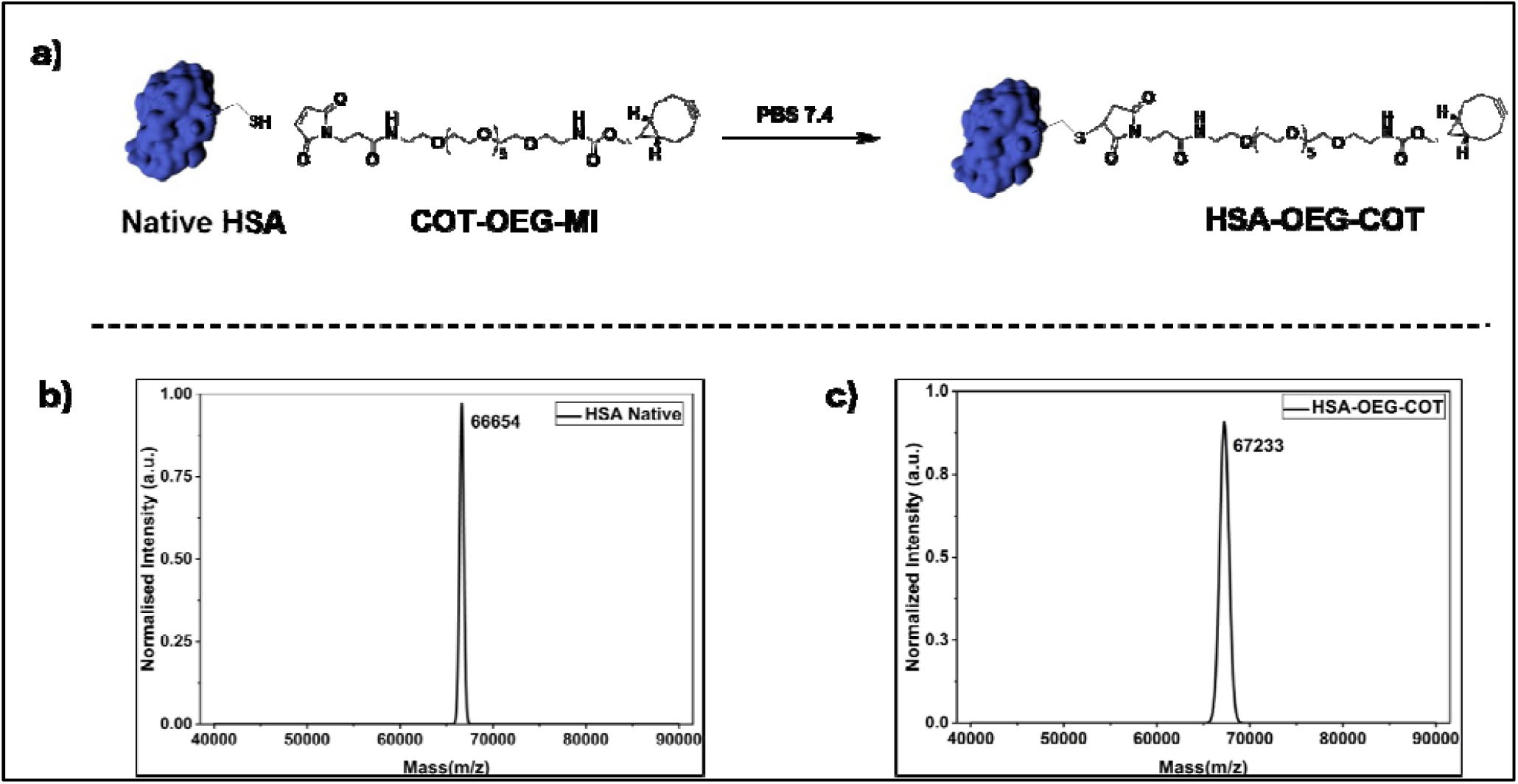
Synthesis of HSA-OEG-COT; a) Schematic representation of HSA-OEG-COT reaction, b) MALDI-TOF MS for native HSA, c) MALDI-TOF MS for HSA-OEG-COT

### Synthesis of facially amphiphilic self-assembling semi-synthetic protein

Following isolation of the HSA-COT conjugate with high purity, strain-promoted azide–alkyne cycloaddition (SPAAC) bioconjugation reaction was carried out under the optimized conditions described in the experimental method section. The reaction progress was monitored directly by MALDI-TOF mass spectrometry without further purification. The mass spectrum displayed a single predominant peak at m/z 68,693 Da corresponding to the expected hydrophilic HSA conjugate, while no signal attributable to native HSA was observed **(Figure 3)**. The observed mass increase was in excellent agreement with the calculated value for the bio-conjugated product, confirming successful bio-orthogonal ligation. The absence of detectable starting material or side products indicates that compound **3f** undergoes highly efficient conjugation with the COT-functionalized protein, affording essentially quantitative conversion under mild aqueous conditions. Having established efficient successful bio-conjugation, the disulfide-linked hydrophilic dendron was selectively removed under reducing conditions to generate the desired self-assembling semi-synthetic protein amphiphile (SSP). Treatment of the purified conjugate with a fivefold molar excess of dithiothreitol (DTT) resulted in rapid and complete cleavage of the disulfide linker. MALDI-TOF mass spectrometric analysis of the crude reaction mixture revealed a single well-resolved peak at m/z 67,874 Da **(Figure 4)**, corresponding to the expected amphiphilic protein conjugate, following quantitative loss of the solubilizing dendron. The absence of additional peaks corresponding to residual conjugated species further confirmed the high purity and structural integrity of the resulting self-assembling semi-synthetic protein (SSP), validating the efficiency of the reductive cleavage under the optimized reaction conditions.

**Figure 3.**
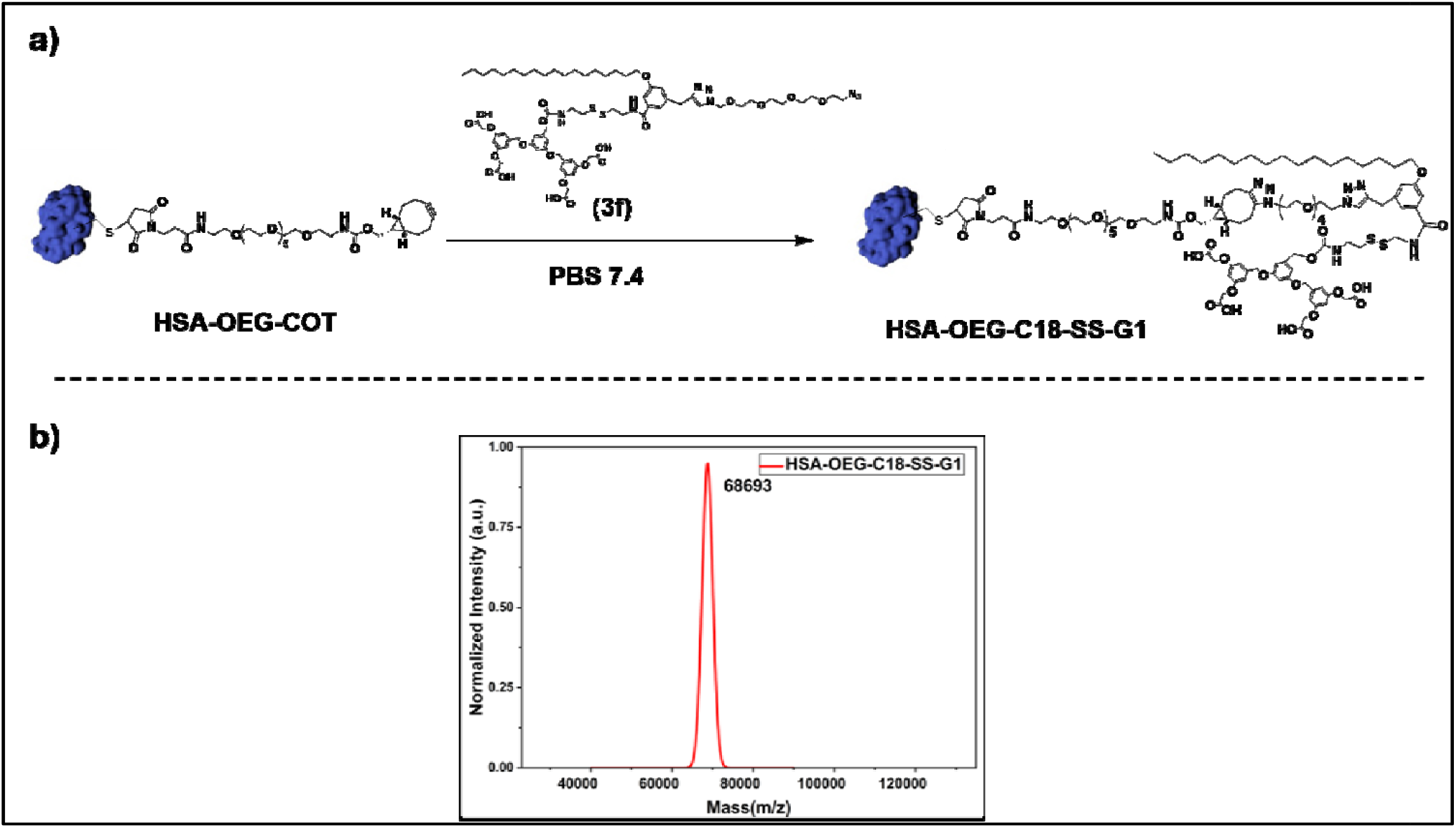
Synthesis of hydrophilic monomeric HSA conjugate; a) Schematic representation of HSA-COT-C18-SS-G1 reaction, b) MALDI-TOF MS for HSA-COT-C18-SS-G1

**Figure 4.**
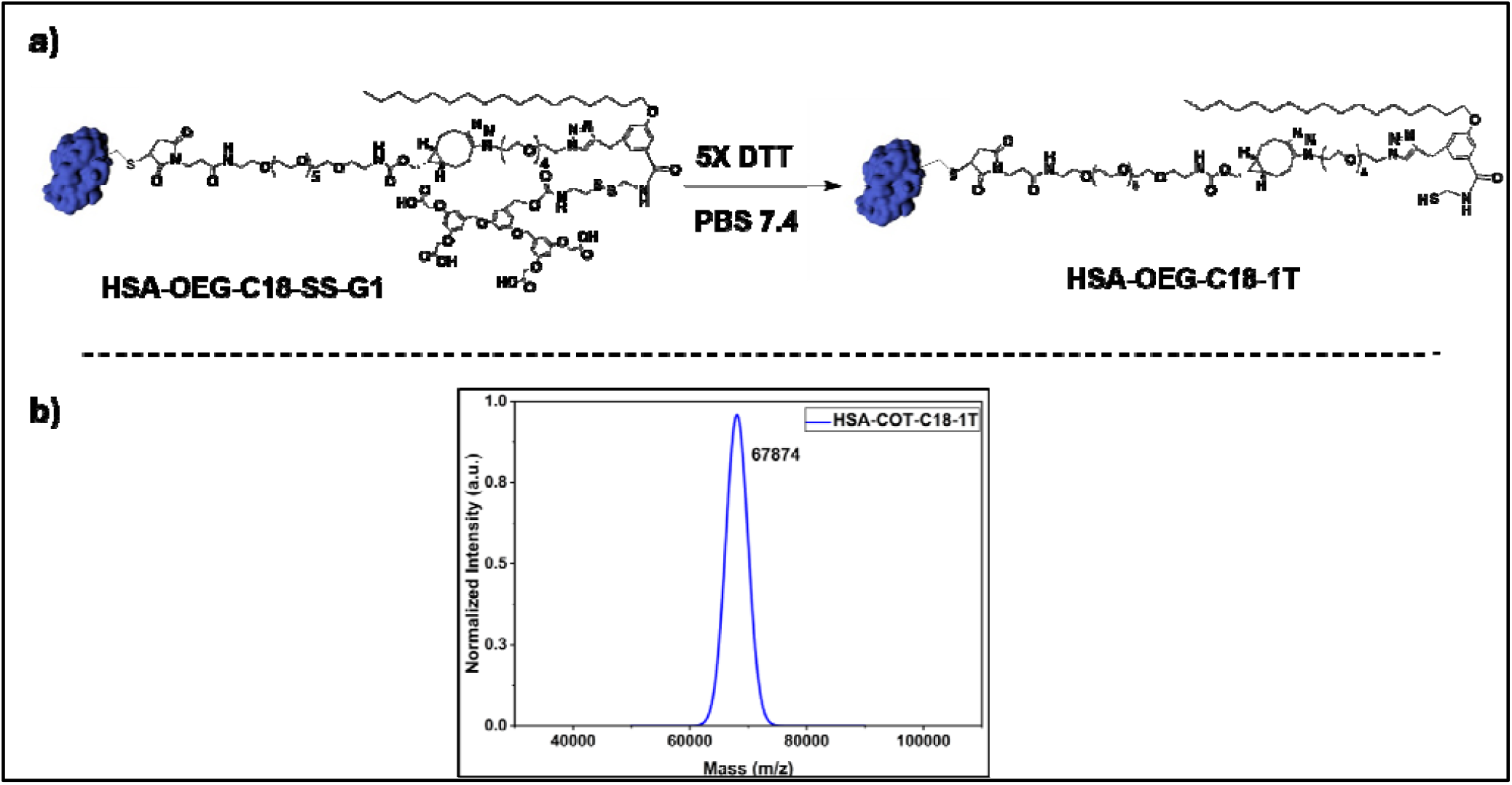
Synthesis of facially amphiphilic SSP; a) Schematic representation of HSA-OEG-C18-1T reaction, b) MALDI-TOF MS for HSA-OEG-C18-1T (m/z 67874)

## Discussion

The chemical synthesis of well-defined, monodisperse SSPs creates opportunities for generating functional biomaterials. The most demanding aspect of this synthesis is the site-specific attachment of a hydrophobic tag to a globular, hydrophilic protein to produce an amphiphilic SSP. Over the years, both MAPLabTech^15–22^ and SAPLabTech^23^ have proven to be robust technologies for preparing functional SSPs with exceptional precision. However, both technologies have certain limitations. Therefore, we sought to develop a complementary chemical method to make SSPs.

Judicious selection of the protein, linker, and hydrophobic moiety is essential for successful molecular design. In this work, our design builds on an earlier report describing an SSP prepared via MAPLabTech: human serum albumin (HSA) as the protein of choice, an oligo (ethylene glycol) (OEG) linker, and an octadecyl group as the hydrophobic moiety.^19^ Synthesizing a chemical probe bearing a solubility tag is a demanding task, so we adopted a convergent strategy to assemble the target chemical probe. Reaction of the target chemical probe with HSA afforded a hydrophilic, water-soluble monomeric HSA bioconjugate. Further, selective removal of the dendritic solubility tag yielded SSP in quantitative yield.

The hydrophilic-lipophilic balance governs the self-assembling behavior of custom-designed proteins.^15^ In this work, a highly hydrophilic dendritic tag is deliberately linked to a water-insoluble hydrophobic chemical probe through a readily cleavable disulfide bond. This design converted the hydrophobic chemical probe into a hydrophilic one and enabled bioconjugation in 100% aqueous medium, producing a hydrophilic monomeric semi-synthetic protein. Following bioconjugation, selective cleavage of the dendritic tag generated a facially amphiphilic SSP.

## Conclusion

In summary, we have disclosed a new chemical technology for preparing well-defined, monodisperse SSPs. This approach employs covalent tagging of a hydrophilic dendritic domain onto a hydrophobic chemical probe to yield a hydrophilic chemical probe that, upon bioconjugation, produces a facially amphiphilic SSP. This technology further expands the chemical toolbox available for generating functional SSPs with applications across biomedical science.

**Scheme 1:**
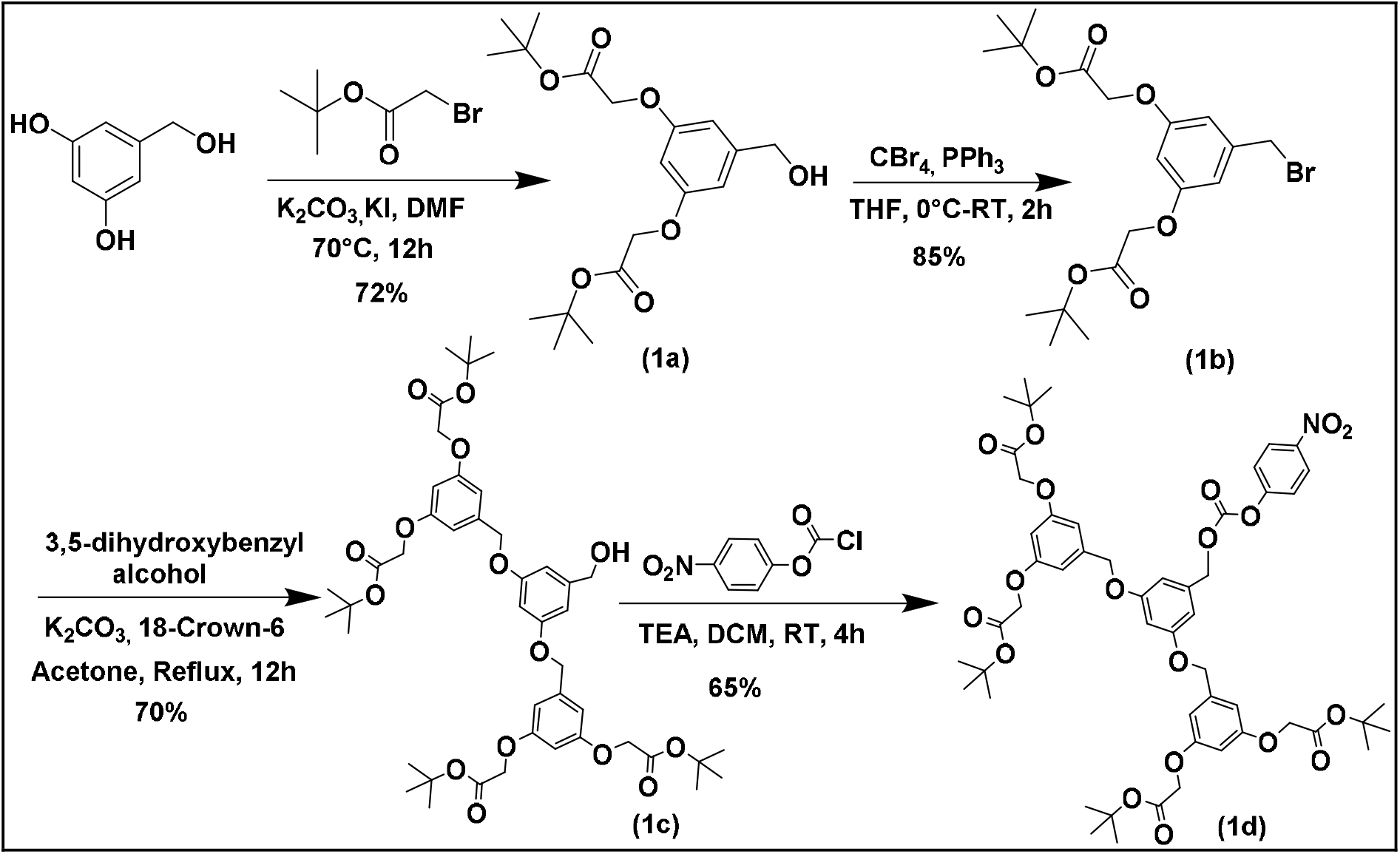
Synthetic scheme for activated dendritic ester

**Scheme 2:**
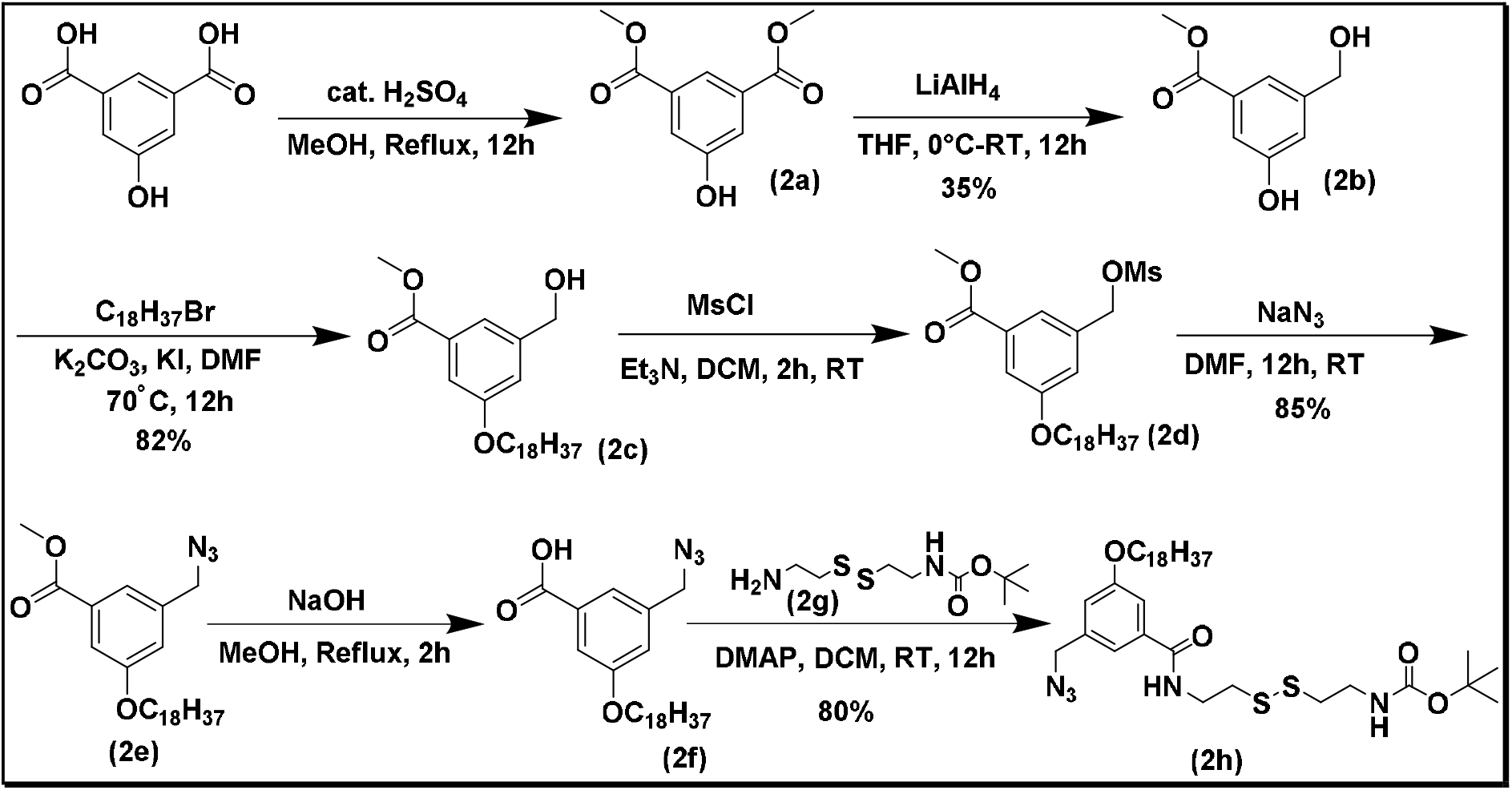
Synthetic scheme for azide-terminated, disulfide-functionalized hydrophobic chemical probe

**Scheme 3:**
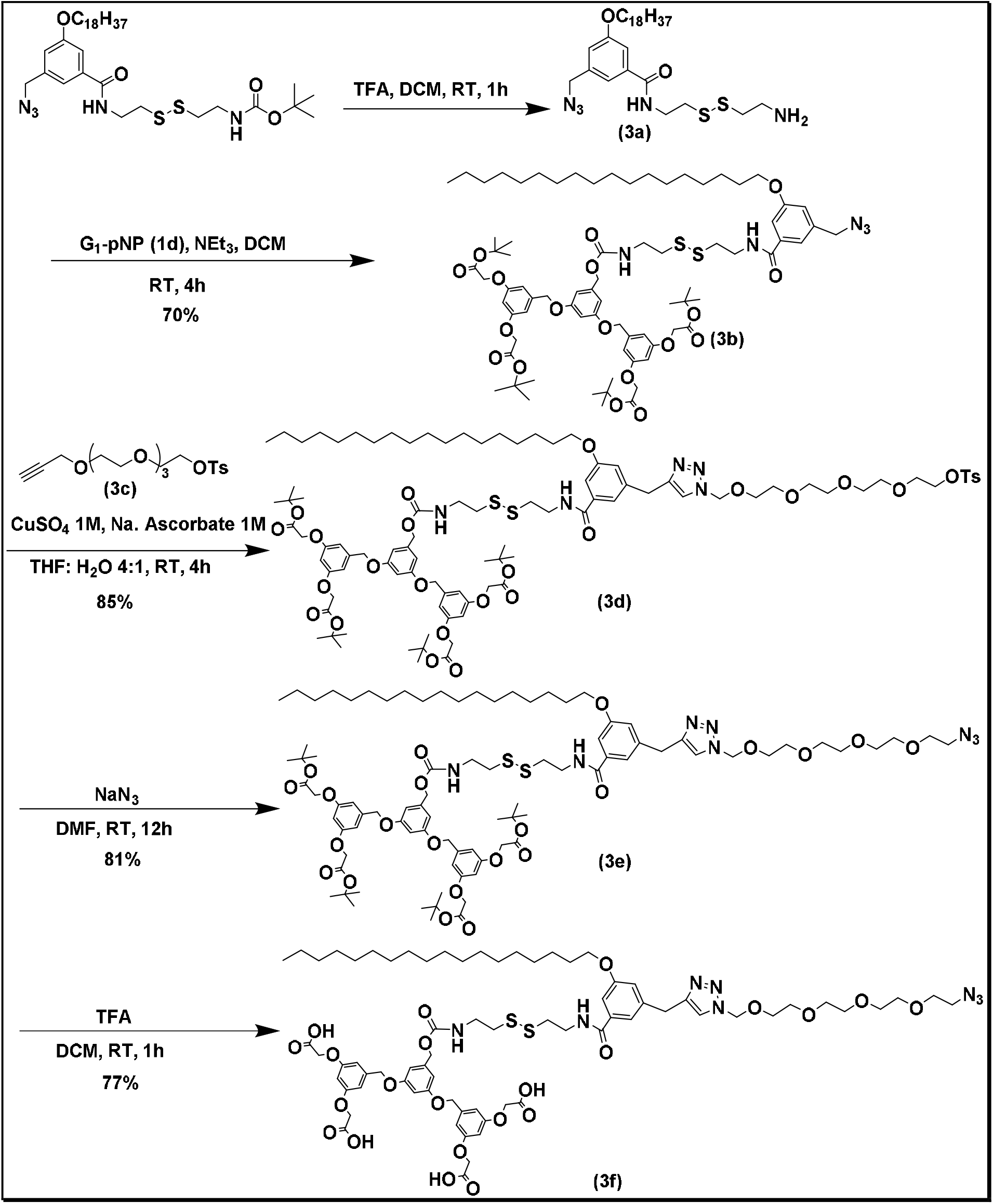
Synthesis of an azide-terminated, disulfide-functionalized hydrophilic chemical probe

## Experimental Section

### General Information

Unless otherwise noted, all reagents and solvents were purchased from commercial suppliers and used as received without further purification. Anhydrous solvents were obtained from commercial sources and handled under standard laboratory conditions where required. High-purity deuterated solvents (CDCl_3_ and D_2_O) were purchased from Sigma-Aldrich and used for NMR analysis.

### Nuclear Magnetic Resonance Spectroscopy

^1^H and ^13^C NMR spectra were recorded on a Bruker 400 MHz spectrometer at ambient temperature. Chemical shifts (δ) are reported in parts per million (ppm) relative to the residual solvent signals of CDCl_3_ (δH 7.26, δC 77.16). Coupling constants (*J*) are reported in hertz (Hz), and signal multiplicities are designated as s (singlet), d (doublet), t (triplet), q (quartet), m (multiplet), and br (broad).

### Protein Selection

Human serum albumin (HSA, 66.6 kDa) was selected as model cysteine-containing proteins for thiol–maleimide conjugation studies. Protein HSA was purchased from Sigma-Aldrich and used without further purification. The molecular weights of the proteins were verified by MALDI-TOF mass spectrometry prior to conjugation.

### MALDI-TOF Mass Spectrometry

For molecular weight determination of HSA protein (>40 kDa), sinapinic acid (15 mg) was dissolved in 1 mL of a 70:30 (v/v) H_2_O/CH_3_CN solution containing 0.1% (v/v) trifluoroacetic acid (TFA) to prepare the matrix solution. Protein samples (50 μM) were mixed with the matrix solution in a 1:5-1:10 (v/v) ratio, and 2 μL of the resulting mixture was spotted onto a MALDI target plate. The spots were allowed to air-dry at room temperature for approximately 30 minutes before analysis. MALDI-TOF mass spectra were acquired over the mass range of m/z 40,000– 80,000 Da.

### Synthesis of HSA-COT

Native human serum albumin (HSA, final concentration 100 μM; theoretical m/z 66,644 Da) was site-selectively modified at the free Cys34 residue using a 2-fold molar excess of a cyclooctyne (COT)-maleimide probe in 50 mM sodium phosphate buffer (pH 7.4) containing 2% (v/v) DMSO. The reaction mixture was incubated at 25°C for 2 hours under gentle end-over-end rotation (20 rpm).

### Sample Preparation for MALDI-TOF MS

3 μL of reaction mixture was mixed with 15 μL of matrix, vortexed, and applied to the MALDI plate. MALDI-TOF MS analysis allowed accurate monitoring of protein modification and confirmation of conjugate formation.

### Synthesis of facially amphiphilic self-assembling semi-synthetic protein

Compound **3f** (2 mg) was dissolved in dimethyl sulfoxide (DMSO, 20 μL) to prepare a 25 mM stock solution. For the conjugation reaction, 2 μL of the stock solution was added to a solution of the protein–COT conjugate (100 μM) prepared in phosphate-buffered saline (PBS, pH 7.4). The total reaction volume was adjusted to 100 μL, corresponding to a final concentration of 500 μM for **3f** (5 eq. relative to the protein) while maintaining the DMSO content at 2% (v/v). The reaction mixture was gently mixed and incubated at 25°C with continuous agitation (20 rpm) for 24 hours to facilitate strain-promoted azide–alkyne cycloaddition (SPAAC)-mediated conjugation. Following incubation, the extent of protein modification was assessed directly by matrix-assisted laser desorption/ionization time-of-flight mass spectrometry (MALDI-TOF MS), which confirmed successful conjugation of **3f** to the protein scaffold. To remove the disulfide-linked solubilizing dendron, dithiothreitol (DTT) was added directly to the reaction mixture to a final concentration of 2.5 mM (5 eq. relative to **3f**). The reduction reaction was allowed to proceed for 1 h at 25°C under gentle agitation (20 rpm). Upon completion, cleavage of the disulfide linker generated the corresponding deprotected protein conjugate, and the reaction progress was subsequently monitored by MALDI-TOF MS.

