## Supplementary material for "Total Synthesis of Self-Assembling Semi-Synthetic Proteins Utilizing a Dendritic Solubility Tag": Supp Info

### Table of Contents

**1. Synthesis of chemical probes and their intermediates**

**2.1. General synthetic scheme for activated G1-ester**

**2.2. General synthetic scheme for synthesis of azide-terminated disulfide-functionalized hydrophobic chemical probe**

**2.3. Synthesis of an azide-terminated, disulfide-functionalized hydrophilic chemical probe**

**3. References**

**4. NMR data**

#### 1. Synthesis of chemical probes and their intermediates

All compounds were characterized using Nuclear Magnetic Resonance Spectroscopy (NMR), specifically $^{1}H$, $^{13}C$. Additionally, they were analysed using either Matrix-Assisted Laser Desorption/Ionization-Time of Flight Mass Spectrometry (MALDI-TOF MS) or High-Resolution Mass Spectrometry (HRMS). The NMR spectra were recorded on a Bruker 400 MHz, with tetramethyl silane (TMS) serving as an internal standard (chemical shifts in ppm). Unless otherwise noted, all $^{1}H$ values are expressed in parts per million (ppm) and are referenced to the residual proton resonances of chloroform ($CHCl_3$) at 7.26 ppm and dimethyl sulfoxide (DMSO) at 2.50 ppm in its respective deuterated solvent. All $^{13}C$ NMR spectra were measured decoupled from the $^{1}H$ nuclei and are also reported in parts per million (ppm).

All reactions were monitored using thin-layer chromatography on precoated silica gel plates, with detection through an ultraviolet (UV) lamp, phosphomolybdic acid (PMA), or ninhydrin staining. Chromatographic separations were carried out using column chromatography with normal-phase silica gel of 100-200 mesh size. The room temperature (RT) ranged from 21 °C to 35 °C. To describe the multiplicities of the signals, the following abbreviations were used: s = singlet, d = doublet, dd = doublet of doublet, t = triplet, q = quartet, m = multiplet, and quint = quintet.

##### 2.1. General synthetic scheme for activated G1-ester

In an oven-dried round-bottomed flask (RBF) containing, 3,5-dihydroxy benzyl alcohol (1.0 eq.), $K_2CO_3$ (2.4 eq.), and tert-butyl bromo acetate (2.2 eq.) taken in DMF. The resulting mixture was heated at 70 °C for 12 hours using an oil bath as a heating source. Upon completion, the reaction mixture was cooled to RT. Then to the reaction mixture, water was added and extracted with ethyl acetate thrice. The combined organic layer was dried over $Na_2SO_4$ and concentrated under reduced pressure to get the crude product, which was purified using Normal-Phase silica gel column chromatography (NPC) with EtOAc/PE as eluent. In an oven-dried RBF, above alcohol (1.0 eq.) and tetra-bromomethane (1.3 eq.) were taken and dissolved in DCM. Then, the solution of triphenylphosphine (1.3 eq.) in DCM was added dropwise at 0 °C and allowed to stir for 10 min at RT. Upon completion of the reaction, water was added to the residue and extracted to the reaction mixture and extracted with DCM. The combined organic layer was dried over $Na_2SO_4$ and concentrated under vacuum to get a crude product, which was purified by silica gel column chromatography using ethyl acetate/hexane. In an oven-dried round-bottomed flask (RBF) containing, 2,3-dihydroxy alcohol (1.0 eq.),

$K_2CO_3$ (2.4 eq.), and above obtained bromide (2.2 eq.) and a catalytic amount of crown ether were dissolved in acetone.

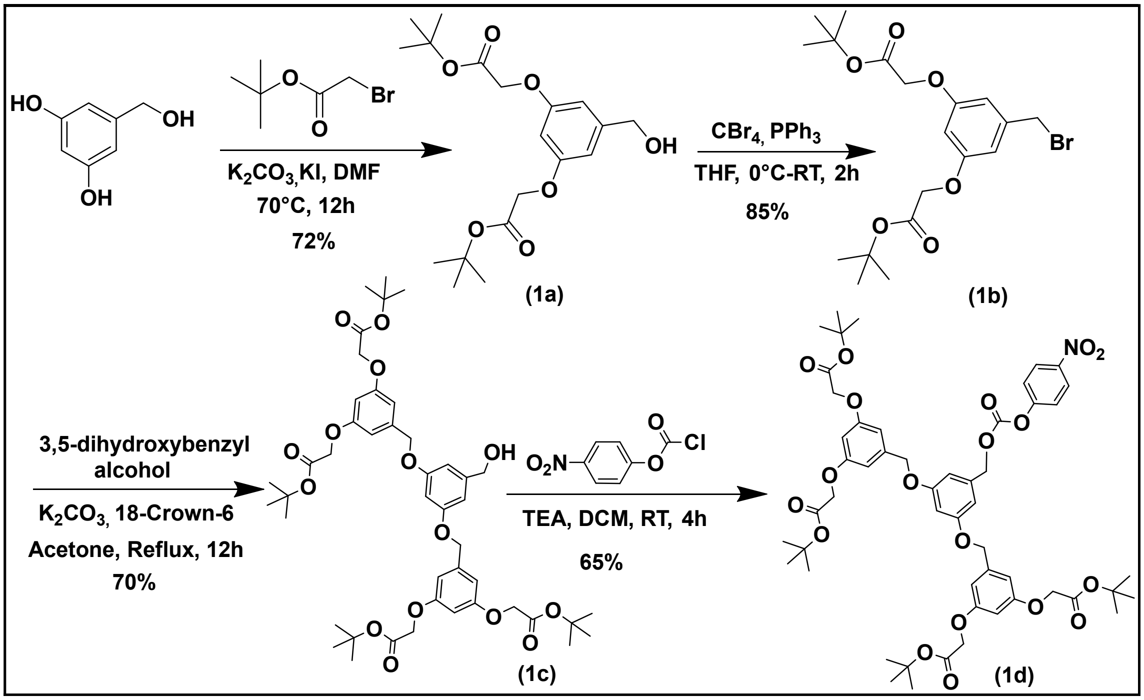


**Scheme 1:** Synthetic scheme for activated G1-ester

The resulting mixture was refluxed for 12 hours using an oil bath as a heating source. Upon completion, the reaction mixture was cooled to RT. Then, acetone was evaporated under reduced pressure. To the obtained residue, water was added and extracted with ethyl acetate thrice. The combined organic layer was dried over $Na_2SO_4$ and concentrated under reduced pressure to get the crude product, which was purified using silica gel column chromatography with EtOAc/PE as eluent. To the above obtained alcohol (1 eq.) taken in RBF, DCM was added. Subsequently para nitro chloroformate (2 eq.) added and kept for stirring. After 10 minutes of stirring, triethylamine (3eq.) added dropwise and allowed to stir 6 hours at RT. Upon completion of reaction, the solvent was evaporated under reduced pressure. To the obtained residue, water was added and extracted with ethyl acetate thrice. The combined organic layer was dried over $Na_2SO_4$ and concentrated under reduced pressure to get the crude product, which was purified using silica gel column chromatography with EtOAc/PE as eluent.

**Characterization of compounds:**

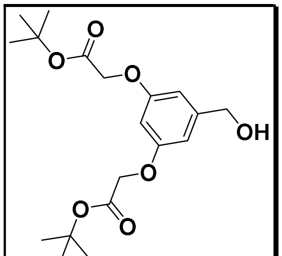

(1a)

**Mol. formula:** $C_{19}H_{28}O_7$

**Mol. weight:** 368.43 g/mol.

**Physical appearance:** Colourless liquid

**Yield:** 72 %

An oven-dried round-bottomed flask was charged with the 2,3-dihydroxy alcohol (3g, 21.41mmol, 1eq.), anhydrous potassium carbonate ($K_2CO_3$, 9.19g, 47.10mmol, 2.2 eq.) and catalytic amount of potassium iodide (KI, 710.74mg, 4.28mmol. 0.2eq.). The reactants were suspended in anhydrous DMF, followed by the dropwise addition of *tert*-butyl bromoacetate (7.4g, 53.52mmol, 2.5 eq.) via syringe at room temperature. The stirred reaction mixture was heated at 70 °C in an oil bath for 12 hours under an inert nitrogen atmosphere. Upon complete consumption of the diol (monitored by TLC), the mixture was cooled to ambient temperature. The crude residue was diluted with distilled water and extracted thoroughly with ethyl acetate (EtOAc) thrice. The combined organic phases were washed with brine, dried over anhydrous sodium sulfate ($Na_2SO_4$), filtered, and concentrated under reduced pressure. Purification of the crude material via silica gel column chromatography using a gradient elution of ethyl acetate in petroleum ether (EtOAc/PE) afforded the compound as a colorless liquid (5.68g, 15.42mmol, 72%), $\mathbf{R_f}$ = 0.65 in 25% EtOAc/PE.

**$^1$H NMR (400 MHz, CDCl$_3$) δ** 6.46 (d, $J$ = 2.3 Hz, 2H), 6.33 (s, 1H), 4.50 (d, $J$ = 5.0 Hz, 2H), 4.41 (s, 4H), 1.43 (s, 18H).

**$^{13}$C NMR (100 MHz, CDCl$_3$) δ** 167.64, 158.74, 143.61, 105.49, 100.55, 82.07, 65.32, 64.38, 27.69.

**MALDI−TOF MS: M+K** 407.11 g/mol.

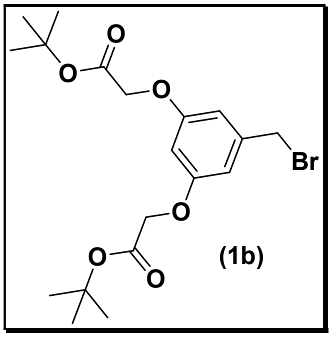

**Mol. formula:** $C_{19}H_{27}BrO_6$

**Mol. weight:** 431.32 g/mol.

**Physical appearance:** Colorless liquid

**Yield:** 85 %

An oven-dried round-bottomed flask was charged with the alcohol intermediate **1a** (5g, 13.57mmol, 1.0 eq.) and tetra-bromomethane ($CBr_4$, 6.75g, 20.36mmol, 1.5eq.) dissolved in anhydrous dichloromethane (DCM). The solution was cooled to 0 °C in an ice bath. Separately, a solution of triphenylphosphine ($PPh_3$, 5.34g, 20.36mmol, 1.5eq.) in anhydrous DCM was prepared and added dropwise to the reaction flask over several minutes. The reaction mixture was maintained at 0 °C for the duration of the addition and then allowed to warm to ambient temperature, where it was stirred for an additional 10 minutes. Upon complete consumption of the starting material (monitored via TLC), the reaction was quenched by the addition of distilled water. The mixture was extracted with DCM thrice. The combined organic phases were washed with brine, dried over anhydrous sodium sulfate ($Na_2SO_4$), filtered, and concentrated under reduced pressure. The resulting crude residue was purified via silica gel column chromatography using a gradient elution of ethyl acetate in hexanes (EtOAc/hexanes) to yield the desired alkyl bromide as colorless liquid (4.97g, 11.52mmol, 85%), $\mathbf{R_f}$ = 0.6 in 20% EtOAc/PE.

**$^1$H NMR (400 MHz, $CDCl_3$) δ** 6.52 (d, *J* = 2.2 Hz, 2H), 6.38 (t, *J* = 2.3 Hz, 1H), 4.46 (s, 4H), 4.35 (s, 2H), 1.46 (s, 18H).

**$^{13}$C NMR (100 MHz, $CDCl_3$) δ** 167.89, 159.39, 140.13, 108.61, 102.34, 82.73, 82.72, 66.00, 33.48, 28.32.

**MALDI−TOF MS: M+K** 471.03 g/mol.

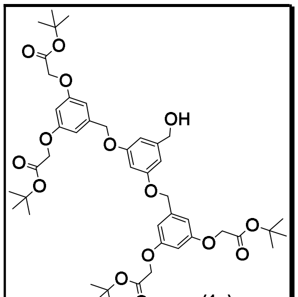

(1c)

Mol. formula: $C_{45}H_{60}O_{15}$

Mol. weight: 840.96 g/mol.

Physical appearance: Colorless liquid

Yield: 70 %

An oven-dried round-bottomed flask was charged with the 3,5-dihydroxybenzyl alcohol (590mg, 4.22mmol, 1.0 eq.), anhydrous potassium carbonate ($K_2CO_3$, 1.4g, 10.12mmol, 2.4 eq.), and a catalytic amount of 18-crown-6 (116mg, 0.42mmol, 0.1 eq.). The mixture was suspended in anhydrous acetone, followed by the addition of the previously synthesized alkyl bromide **1b** (4g, 9.27mmol, 2.2 eq.). The flask was fitted with a reflux condenser, and the stirred reaction mixture was heated at reflux (56 °C) in an oil bath for 12 hours under an inert nitrogen atmosphere. Upon complete consumption of the starting material (monitored by TLC), the mixture was cooled to ambient temperature, and the volatile components were evaporated under reduced pressure. The remaining residue was partitioned between distilled water and ethyl acetate (EtOAc). The layers were separated, and the aqueous phase was extracted with EtOAc. The combined organic layers were washed with brine, dried over anhydrous sodium sulfate ($Na_2SO_4$), filtered, and concentrated under reduced pressure. The resulting crude residue was purified via silica gel column chromatography using a gradient elution of ethyl acetate in petroleum ether (EtOAc/PE) to afford the compound as colorless liquid **1c** (2.48g, 2.95mmol) as 70% yield. $\mathbf{R_f}$ = 0.65, solvent 50% (EtOAc/PE)

**$^1$H NMR (400 MHz, $CDCl_3$) δ** 6.56 (dd, $J$ = 4.1, 2.3 Hz, 6H), 6.47 – 6.38 (m, 3H), 4.91 (s, 4H), 4.46 (s, 8H), 1.46 (s, 36H).

**$^{13}$C NMR (100 MHz, $CDCl_3$) δ** 167.51, 159.59, 158.88, 143.49, 139.25, 106.13, 105.38, 101.06, 82.13, 69.36, 65.42, 64.70, 27.72.

**MALDI−TOF MS: M+K** 879.34 g/mol.

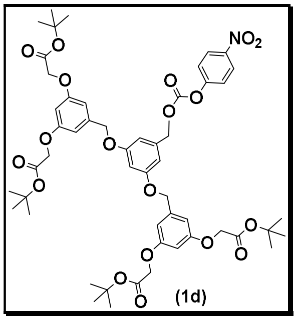


**Mol. formula:** $C_{52}H_{63}NO_{19}$

**Mol. weight:** 1006.06 g/mol.

**Physical appearance:** Colorless liquid

**Yield:** 65 %

To a stirred solution of the alcohol intermediate **1c** (2g, 2.38mmol, 1.0 eq.) in anhydrous dichloromethane (DCM) in a round-bottomed flask, *p*-nitrophenyl chloroformate (958mg, 4.76mmol, 2.0 eq.) was added in one portion at room temperature. The mixture was allowed to stir for 10 minutes to ensure full dissolution and equilibrium. Subsequently, triethylamine ($Et_3N$, 1.22ml, 9.5mmol 4.0 eq.) was added dropwise via syringe. The resulting reaction mixture was allowed to stir at ambient temperature for 6 hours. Upon complete consumption of the starting material (monitored by TLC), the volatile components were removed under reduced pressure. The remaining crude residue was partitioned between distilled water and ethyl acetate (EtOAc). The layers were separated, and the aqueous phase was extracted thoroughly with EtOAc thrice. The combined organic layers were washed with brine, dried over anhydrous sodium sulfate ($Na_2SO_4$), filtered, and concentrated under reduced pressure. Purification of the crude material via silica gel column chromatography using a gradient elution of ethyl acetate in petroleum ether (EtOAc/PE) afforded the pure compound **1d** as colorless liquid (1.56g, 2.36mmol) with 65.2% yield. **$R_f$** 0.5, solvent 30% (EtOAc/PE).

**$^1$H NMR (400 MHz, $CDCl_3$) δ:** 8.27 (d, $J$ = 9.2 Hz, 2H), 7.38 (d, $J$ = 9.1 Hz, 2H), 6.67 – 6.53 (m, 7H), 6.43 (t, $J$ = 2.3 Hz, 2H), 5.20 (s, 2H), 4.95 (s, 4H), 4.48 (s, 8H), 1.47 (s, 36H).

**$^{13}$C NMR (100 MHz, $CDCl_3$) δ:** 167.74, 160.07, 159.27, 139.22, 136.47, 125.31, 121.81, 107.47, 106.51, 102.40, 101.39, 82.47, 70.69, 69.84, 65.76, 28.05.

**MALDI TOF MS: M+K** 1028.34 g/mol.

#### 2.2. General synthetic scheme for synthesis of azide-terminated disulfide-functionalized hydrophobic chemical probe

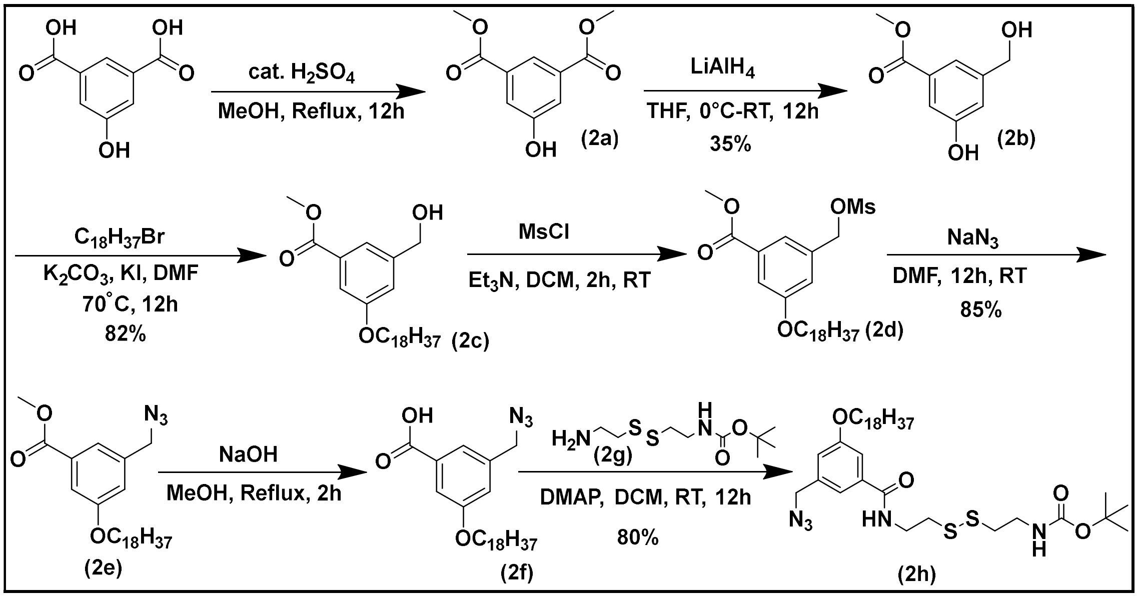


**Scheme 2:** Synthetic scheme of azide-terminated disulfide-functionalized hydrophobic chemical probe

To a stirred solution of 5-hydroxyisophthalic acid (1eq.) in anhydrous methanol (30 mL) at room temperature, concentrated sulfuric acid (3eq.) was added dropwise. The reaction mixture was heated under reflux for 12 hours, cooled to room temperature, and concentrated *in vacuo*. The resulting residue was dissolved in ethyl acetate (100 mL), washed sequentially with saturated aqueous $NaHCO_3$, brine and dried over $Na_2SO_4$, filtered, and concentrated under reduced pressure to afford dimethyl 5-hydroxyisophthalate **(2a)** as a white solid. To a stirred ice-cold solution of dimethyl 5-hydroxyisophthalate, (1 eq.) in 140 ml dry THF, $LiAlH_4$ (1M solution in THF, 0.75 eq.) was added dropwise for 30 min. Toward the end of the addition 60 ml THF was added to the suspension to enable efficient stirring. The yellow-orange color of the suspension was changed to white and the suspension was stirred for 1.5 hours at 25 °C. Water was added to quench the excess of $LiAlH_4$ and most of the THF was evaporated. 30 ml 2N HCl was added to destroy the aluminate complexes. Sodium bicarbonate was added to the solution until it reached pH 7. The yellow cloudy solution was filtered and the white solid was discarded. The filtrate was saturated with NaCl and extracted with EtOAc. The aqueous phase was extracted again with EtOAc. The combined organic layers were dried on $MgSO_4$ and evaporated under reduced pressure. The residue was purified by flash chromatography on silica gel using EA/PE as eluent to get product **(2b)**. In an oven-dried round-bottomed flask (RBF) containing, above obtained mono-ester (1.0 eq.), $K_2CO_3$ (2.4 eq.), and 1-bromooctadecane (2.2 eq.) and a catalytic amount of KI taken in DMF. The resulting mixture was heated at 70 °C for 12 hours using an oil bath as a heating source. Upon completion, the reaction mixture was cooled to RT. Then to the reaction mixture, water was added and extracted with ethyl acetate thrice. The combined organic layer was dried over $Na_2SO_4$ and concentrated under reduced pressure to get the crude product, which was purified using silica gel column chromatography with EtOAc/PE as eluent. In an oven-dried round-bottomed flask (RBF), above obtained compound **2c** and MsCl (2eq.) dissolved in DCM. After 10 minutes, 3 eq. of triethyl-amine added dropwise and reaction mixture was kept for stirring. Upon completion of the reaction, solvent and excess triethyl amine evaporated. The residue extracted with ethyl acetate and washed with brine thrice. The organic layer was dried over $Na_2SO_4$ and concentrated under reduced pressure to get the crude product **2d**, which was used for next step without purification. The above crude dissolved in DMF and 10 eq. sodium azide was added and reaction mixture kept for stirring overnight. Upon completion, water was added to the reaction mixture and extracted thrice with DCM. Combined organic layer was dried over $Na_2SO_4$ and concentrated under vacuum to get crude product which was purified using silica gel column chromatography using EtOAc/hexane to obtain **2e** as white solid. To the solution of above obtained azide in THF/MeOH (2:1), aq. solution of KOH (5 eq.) was added. The resultant mixture was refluxed for 12 hours using an oil bath as a heating source. After completion, the solvent was removed under reduced vacuum. To the obtained residue, excess of water was added and the mixture was refluxed for 6 hours. The mixture was cooled to ambient temperature and the aq. solution of HCl (20%) was added dropwise to get a white precipitate, which was filtered and dried under vacuum to get a crude product **2f** which was utilized without further purification. Cystamine dihydrochloride (1eq) was taken in a RBF and MeOH was added. While in stirring triethyl amine (4 eq.) added dropwise until clear solution formed. After 30 minutes of stirring, di-Boc (0.2eq.) added and kept for stirring 2 hours. Upon completion of reaction, solvent was evaporated, to the obtained residue, water was added and extracted with ethyl acetate thrice. The combined organic layer was dried over $Na_2SO_4$ and concentrated under reduced pressure to get the crude product **2g**, and directly used for next step without purification. Acid compound **(2f)** was taken and DCM was added, HATU (2 eq.), DMAP (2 eq.) was added to the solution and kept for stirring. After 15 min, mono-Boc terminated compound **(2g)** (4 eq.) was added and kept for 12 hours at RT. Upon completion, product was extracted thrice with DCM and brine. Combined organic layer was dried over $Na_2SO_4$ and concentrated under vacuum to get crude product which was purified using silica gel column chromatography using EtOAc/hexane.

**<u>Characterization of compounds:</u>**

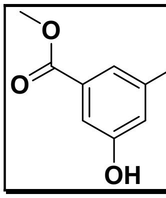

(2a)

**Mol. formula:** $C_{10}H_{10}O_5$

**Mol. weight:** 210.19 g/mol.

**Physical appearance:** White solid

**Yield:** Quantitative

To a stirred solution of 5-hydroxyisophthalic acid (8.00 g, 44 mmol, 1.0 eq.) in anhydrous methanol (120 mL, 0.18 M) in a 250 mL round-bottomed flask was added concentrated sulfuric acid (0.48 mL, 9.6 mmol, 0.22 eq.) dropwise at room temperature. The flask was equipped with a reflux condenser, and the reaction mixture was heated under reflux in an oil bath for 12 hours. The progress of the reaction was monitored by thin-layer chromatography (TLC). Upon complete consumption of the starting material, the mixture was cooled to room temperature, and the volatile solvent was removed *in vacuo* under reduced pressure. The resulting residue was dissolved in ethyl acetate (150 mL) and transferred to a separatory funnel. The organic phase was washed sequentially with saturated aqueous sodium bicarbonate ($NaHCO_3$), and brine thrice. The organic layer was then separated, dried over anhydrous sodium sulfate ($Na_2SO_4$) filtered, and concentrated under reduced pressure to afford the pure dimethyl ester as a white solid (9.15g, quantitative yield, 100%). No further purification was required and used directly for next step.

**$^1$H NMR (400 MHz, DMSO-d$_6$) δ** 10.31 (s, 1H), 7.87 (t, $J$ = 1.5 Hz, 1H), 7.51 (d, $J$ = 1.5 Hz, 2H), 3.83 (s, 6H).

**$^{13}$C NMR (100 MHz, DMSO-d$_6$) δ** 165.73, 158.16, 131.63, 120.63, 120.45, 52.67.

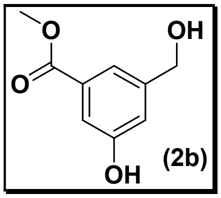

**Mol. formula:** $C_9H_{10}O_4$

**Mol. weight:** 180.14 g/mol.

**Physical appearance:** White solid

**Yield:** 35%

An anhydrous solution of dimethyl 5-hydroxyisophthalate (9 g, 42.82 mmol, 1.0 eq.) in dry tetrahydrofuran (THF, 140 mL) was prepared in a dry, round-bottomed flask under an inert nitrogen atmosphere. The mixture was cooled to 0°C using an ice-water bath and stirred vigorously. Lithium aluminum hydride ($LiAlH_4$) (2.93g, 77.07 mmol, 1.8 eq.) was added dropwise to the cold solution over a period of 2 hours. Due to the formation of a thick suspension as the reaction progressed, an additional portion of dry THF (60 mL) was added toward the end of the addition to maintain efficient magnetic stirring. During this process, the colour of the reaction mixture shifted from a distinct yellow-orange to a characteristic white suspension. After completing the addition, the cooling bath was removed, and the mixture was allowed to warm to room temperature (25 °C) and stirred for an additional 1.5 hours. To quench the reaction, the mixture was cooled back to 0 °C, and water was carefully added dropwise to decompose the excess, unreacted $LiAlH_4$. The majority of the volatile THF solvent was then removed under reduced pressure using a rotary evaporator. To break up the insoluble, aluminum-oxygen complexes, aqueous hydrochloric acid (30 mL of a 2N solution) was added to the residue. The mixture was stirred until the salts dissolved, and then neutralized to pH 7 by the cautious, portion-wise addition of solid sodium bicarbonate $NaHCO_3$. The resulting yellow, cloudy suspension was filtered through a pad of Celite to remove insoluble inorganic byproducts, and the discarded white filter cake was washed thoroughly with ethyl acetate. The biphasic filtrate was saturated with sodium chloride NaCl to minimize the solubility of the product in the aqueous layer, and the organic layer was separated. The remaining aqueous phase was thoroughly back-extracted with ethyl acetate. The combined organic extracts were dried over anhydrous $Na_2SO_4$, filtered to remove the drying agent, and concentrated *in vacuo* under reduced pressure. The crude residue was purified by normal phase column chromatography on silica gel, utilizing a gradient elution of ethyl acetate in hexanes, to afford the desired partially reduced mono-alcohol product as white powder (2.74g, 15.04mmol) with 35% yield. **$R_f$** 0.5 in 25% EA/PE.

**$^{1}$H NMR (400 MHz, DMSO-$d_6$)** δ 9.77 (s, 1H), 7.39 (d, *J* = 1.5 Hz, 1H), 7.24 (t, *J* = 2.0 Hz, 1H), 7.02 (dd, *J* = 2.5, 1.4 Hz, 1H), 5.31 (t, *J* = 5.8 Hz, 1H), 4.49 (d, *J* = 5.7 Hz, 2H), 3.81 (s, 3H).

**$^{13}$C NMR (100 MHz, DMSO-$d_6$)** δ 166.83, 157.87, 145.19, 131.06, 118.52, 118.31, 114.50, 62.87, 52.37.

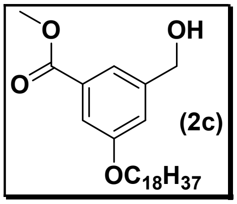


**Mol. formula:** $C_{27}H_{46}O_4$

**Mol. weight:** 434.14 g/mol.

**Physical appearance:** White solid

**Yield:** 82%

An oven-dried round-bottomed flask equipped with a magnetic stir bar was charged with the previously obtained mono-ester intermediate (1.5g, 8.23mmol, 1.0 eq.), anhydrous potassium carbonate ($K_2CO_3$, 16.47mmol, 2 eq.), and 1-bromooctadecane (4.12g, 12.35mmol, 1.5eq.). The mixture was dissolved in DMF to achieve a heterogenous concentration. The flask was heated at 70 °C for 12 hours utilizing a preheated oil bath as the thermal source. Reaction progress and consumption of the starting material were periodically monitored by TLC. Upon quantitative consumption of the mono-ester, the reaction mixture was allowed to cool to ambient temperature. Further it was diluted with water, then transferred to a separatory funnel. The aqueous phase was thoroughly extracted with ethyl acetate thrice. The organic layers were washed with brine, and dried over anhydrous $Na_2SO_4$ filtered to remove the drying agent, and the filtrate was concentrated under reduced pressure. The crude material was subsequently purified via NPC on silica gel, utilizing a gradient elution profile of ethyl acetate in petroleum ether (EtOAc/PE), to yield the pure alkylated target compound as white solid (2.93g, 6.74mmol) with 82% yield. $\mathbf{R_f}$ 0.6 in 20% EA/PE.

**$^1$H NMR (400 MHz, $CDCl_3$) δ** 7.59 (s, 1H), 7.46 (d, *J* = 1.1 Hz, 1H), 7.12 (s, 1H), 4.70 (d, *J* = 4.1 Hz, 2H), 3.99 (s, 2H), 3.90 (s, 3H), 1.92 (s, 1H), 1.78 (dd, *J* = 8.2, 6.4 Hz, 2H), 1.25 (s, 34H), 0.88 (t, *J* = 6.7 Hz, 3H).

**$^{13}$C NMR (100 MHz, $CDCl_3$) δ** 166.77, 159.25, 142.50, 131.33, 119.83, 117.89, 113.82, 68.15, 64.57, 52.01, 31.75, 29.52, 29.51, 29.48, 29.43, 29.41, 29.20, 29.19, 29.01, 25.83, 22.51, 13.94.

**MALDI M+K** 470.4 g/mol.

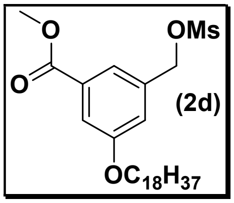


**Mol. formula:** $C_{28}H_{48}O_6S$

**Mol. weight:** 512.32 g/mol

**Physical appearance:** Not determined

**Yield:** Not calculated

An oven-dried round-bottomed flask equipped with a magnetic stir bar was charged with compound **2c** (2.5g, 5.75mmol, 1.0 eq.) and dissolved in anhydrous dichloromethane (DCM) under an inert atmosphere. Methanesulfonyl chloride (MsCl, 1.52ml, 3.0 eq.) was added to the solution, and the mixture was allowed to stir at ambient temperature for 10 minutes to ensure complete homogenization. Subsequently, anhydrous triethylamine ($NEt_3$, 2.49ml, 3.0 eq.) was added dropwise to the reaction mixture. The reaction mixture was maintained under continuous stirring until TLC indicated the complete consumption of the starting material. Upon completion, the volatile solvent (DCM) and excess triethylamine were removed *in vacuo* under reduced pressure using a rotary evaporator. The resulting crude residue was partitioned between ethyl acetate (EtOAc) and water, then transferred to a separatory funnel. The organic layer was isolated and washed sequentially with brine thrice to remove residual ammonium salts. The organic phase was dried over anhydrous sodium sulfate ($Na_2SO_4$), filtered, and concentrated under reduced pressure to afford the crude mesylate target intermediate product which was utilized immediately in the subsequent synthetic step without further chromatographic purification.

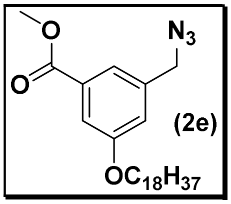


**Mol. formula:** $C_{27}H_{45}N_3O_3$

**Mol. weight:** 459.68 g/mol.

**Physical appearance:** White Solid

**Yield:** 85%

The previously obtained crude mesylate intermediate (2.90g, 5.66mmol, 1.0 eq.) was dissolved in anhydrous *N*,*N*-dimethylformamide (DMF) in a round-bottomed flask equipped with a magnetic stir bar. Sodium azide ($NaN_3$, 3.68g, 56.56mmol, 10.0 eq.) was added in one portion to the solution. The resulting heterogeneous mixture was stirred continuously at ambient temperature overnight. Upon completion, the reaction mixture was quenched by the careful addition of deionized water and transferred to a separatory funnel. The aqueous phase was thoroughly extracted with dichloromethane thrice. The organic layers were combined, washed with water and brine to remove residual DMF, and dried over anhydrous sodium sulfate $Na_2SO_4$. The drying agent was removed via filtration, and the filtrate was concentrated *in vacuo* under reduced pressure. The resulting crude material was purified by NPC on silica gel, utilizing a gradient elution system of ethyl acetate in hexanes (EtOAc/hexane), to afford the pure azide target compound as white solid (2.21g, 4.81mmol) with 85% yield. **$R_f$** 0.8 in 10% EA/PE.

**$^1$H NMR (400 MHz, $CDCl_3$) δ** 7.60 – 7.45 (m, 2H), 7.05 (d, *J* = 2.0 Hz, 1H), 4.35 (s, 2H), 4.00 (t, *J* = 6.5 Hz, 2H), 3.91 (s, 3H), 1.85 – 1.74 (m, 2H), 1.47 – 1.25 (m, 32H), 0.87 (t, *J* = 6.7 Hz, 3H).

**$^{13}$C NMR (100 MHz, $CDCl_3$) δ** 166.40, 159.36, 136.93, 131.69, 121.07, 119.12, 114.41, 68.21, 54.14, 52.07, 31.74, 29.51, 29.47, 29.41, 29.38, 29.18, 28.96, 25.81, 22.50, 13.92.

**MALDI:** **M+Na** 482.6 g/mol.

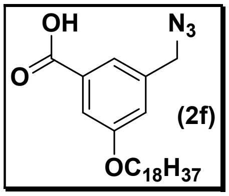

**Mol. formula:** $C_{26}H_{43}N_3O_3$

**Mol. weight:** 445.65 g/mol.

**Physical appearance:** White Solid

**Yield:** Quantitative

To a stirred solution of the previously obtained azide (2.1g, 4.57mmol) in a 2:1 mixture of tetrahydrofuran (THF) and methanol (MeOH), an aqueous solution of sodium hydroxide (NaOH, 0.91g, 22.4mmol, 5.0 eq.) was added. The resulting reaction mixture was heated at reflux in an oil bath for 12 hours. Upon consumption of the starting material, the volatile solvents were removed under reduced pressure. The remaining residue was diluted with an excess of water, and the resulting mixture was acidified by the dropwise addition of a 20% aqueous hydrochloric acid (HCl) solution until a white precipitate formed. The precipitate was collected via filtration and dried under vacuum to afford the crude product, which was utilized in the subsequent step without further purification.

**$^1$H NMR (400 MHz, DMSO-d$_6$) δ** 7.51 (s, 1H), 7.42 – 7.33 (m, 1H), 7.23 – 7.11 (m, 1H), 4.49 (s, 2H), 4.05 – 3.96 (m, 3H), 1.71 (p, *J* = 6.8 Hz, 2H), 1.22 (s, 32H), 0.84 (t, *J* = 6.4 Hz, 3H).

**$^{13}$C NMR (100 MHz, DMSO-d$_6$) δ** 138.19, 121.73, 119.56, 114.72, 68.32, 53.52, 31.80, 29.49, 29.20, 29.01, 25.91, 22.60, 14.45.

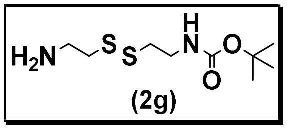

Mol. formula: $C_9H_{20}N_2O_2S_2$

Mol. weight: 252.39 g/mol.

Physical appearance: colorless liquid

Yield: 42%

To a stirred suspension of cystamine dihydrochloride (5g, 32.84mmol, 1.0 eq.) in methanol (MeOH) in a round-bottom flask, triethylamine ($Et_3N$, 6.73ml, 130.52mmol, 4.0 eq.) was added dropwise at ambient temperature until a homogeneous solution was obtained. After stirring for 30 minutes, di-*tert*-butyl dicarbonate ($Boc_2O$, 2.87g, 13.13mmol, 0.4 eq.) was added, and the reaction mixture was allowed to stir for an additional 2 hours. Upon completion, the volatile components were removed under reduced pressure. The resulting residue was diluted with water and extracted with ethyl acetate (EtOAc) thrice. The combined organic layers were dried over anhydrous sodium sulfate $Na_2SO_4$, filtered, and concentrated under reduced pressure. The crude material was purified via NPC, eluting with a gradient of methanol in dichloromethane (MeOH/DCM), to afford the desired product as colorless liquid (1.4g, 5.55mmol) with 42% yield. **$R_f$** 0.2 in (5% MeOH/DCM+1% TEA).

**$^1$H NMR (400 MHz, $CDCl_3$) δ** 3.41 (d, $J$ = 6.2 Hz, 2H), 2.97 (td, $J$ = 6.3, 1.9 Hz, 2H), 2.74 (q, $J$ = 6.7 Hz, 4H), 1.56 – 1.48 (m, 2H), 1.40 (s, 9H).

**$^{13}$C NMR (100 MHz, $CDCl_3$) δ** 156.16, 79.84, 42.78, 40.90, 39.65, 38.74, 28.76.

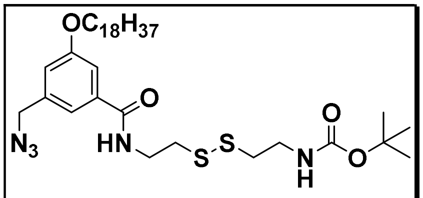


(2h)

**Mol. formula:** $C_{35}H_{61}N_5O_4S_2$

**Mol. weight:** 680.02 g/mol.

**Physical appearance:** White solid

**Yield:** 80%

To a stirred solution of carboxylic acid (0.8g, 1.8mmol, 1.0 eq.) in anhydrous dichloromethane (DCM), 1-[bis(dimethylamino)methylene]-1H-1,2,3-triazolo[4,5-b]pyridinium 3-oxide hexafluorophosphate (HATU, 1.37g, 3.59mmol, 2.0 eq.) and 4-dimethylaminopyridine

(DMAP, 678mg, 5.39mmol, 3.0 eq.) were added sequentially at room temperature. The mixture was allowed to stir for 15 minutes to facilitate in situ carboxylate activation. Subsequently, a solution of the mono-Boc-protected amine (1.36g, 5.39mmol, 3.0 eq.) in DCM was added dropwise. The resulting reaction mixture was stirred at ambient temperature for 6 hours. Upon consumption of the starting material (monitored by TLC), the mixture was diluted with DCM and washed sequentially with water and brine. The layers were separated, and the organic phase was dried over anhydrous sodium sulfate ($Na_2SO_4$), filtered, and concentrated under reduced pressure. The crude residue was purified via NPC using an ethyl acetate/hexane gradient (EtOAc/hexane) to yield the desired product as white solid (980mg, 1.44mmol) with 80% yield. $\mathbf{R_f}$ 0.6 at 35% EA/PE.

**$^1$H NMR (401 MHz, $CDCl_3$) δ** 7.31 (d, *J* = 2.1 Hz, 1H), 7.01 – 6.93 (m, 2H), 5.01 (t, *J* = 6.1 Hz, 1H), 4.34 (s, 2H), 3.98 (t, *J* = 6.6 Hz, 2H), 3.76 (d, *J* = 6.1 Hz, 2H), 3.45 (d, *J* = 6.6 Hz, 2H), 2.94 (s, 2H), 2.81 (t, *J* = 6.5 Hz, 2H), 1.78 (d, *J* = 6.2 Hz, 3H), 1.41 (s, 9H), 1.25 (s, 30H), 0.87 (t, *J* = 6.6 Hz, 3H).

**$^{13}$C NMR (100 MHz, $CDCl_3$) δ** 166.97, 159.52, 155.70, 137.07, 136.05, 118.20, 117.23, 112.71, 79.44, 68.18, 54.19, 39.19, 38.78, 37.99, 37.89, 31.71, 29.49, 29.47, 29.44, 29.40, 29.37, 29.18, 29.15, 28.96, 28.16, 25.80, 22.48.

**MALDI: M+Na** 703.2 g/mol.

##### 2.3. Synthesis of an azide-terminated, disulfide-functionalized hydrophilic chemical probe

To a stirred solution of compound **2h** (1.0 eq.) in anhydrous dichloromethane (DCM) taken in an oven-dried round-bottom flask, trifluoroacetic acid (TFA, 3.0 eq.) was added dropwise at room temperature. The reaction mixture was stirred at ambient temperature for 2 hours. Upon complete consumption of the starting material (monitored by TLC), the mixture was cooled to 0°C and quenched via the careful addition of a saturated aqueous sodium bicarbonate ($NaHCO_3$) solution until the aqueous layer reached pH 8. The organic layer was separated, and the aqueous phase was extracted with DCM. The combined organic layers were dried over anhydrous sodium sulfate ($Na_2SO_4$), filtered, and concentrated under reduced pressure to afford the crude amine intermediate **3a**, which was utilized immediately in the subsequent step without further purification.

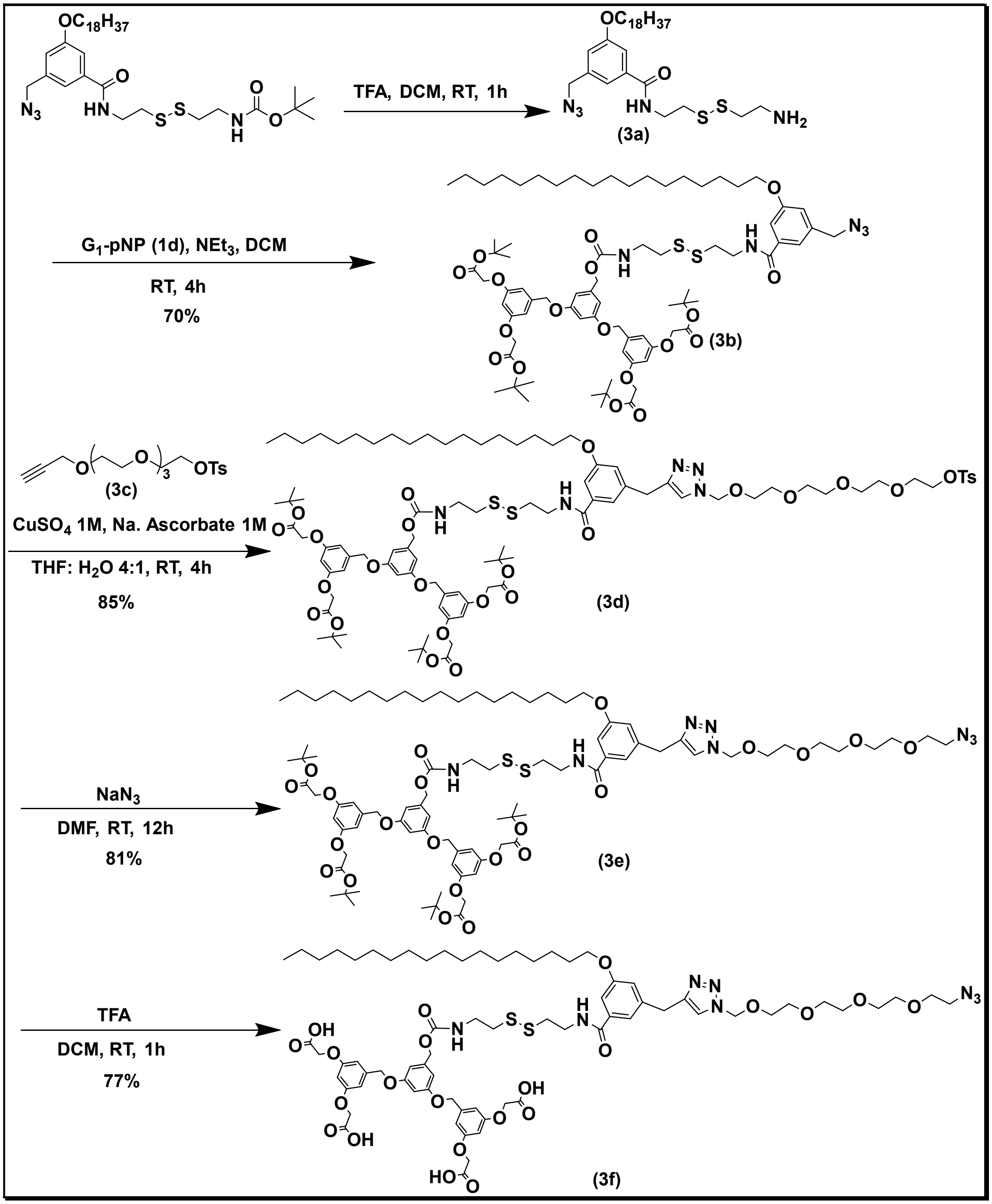


**Scheme 3:** Synthetic scheme of an azide-terminated, disulfide-functionalized hydrophilic chemical probe

The crude intermediate was dissolved in anhydrous DCM, and compound (1.0 eq.) was added to the solution. The mixture was stirred at room temperature for 10 minutes, followed by the dropwise addition of triethylamine ($Et_3N$, 2.0 eq.). The resulting reaction mixture was allowed to stir for 4 hours. The mixture was then diluted with water, and the organic layer was separated.

The organic phase was washed sequentially with saturated aqueous $NaHCO_3$ and brine, dried over anhydrous $Na_2SO_4$, filtered, and concentrated under reduced pressure. Purification of the crude residue via silica gel column chromatography using an ethyl acetate/hexane gradient (EtOAc/hexane) afforded the desired compound **3b**. The azide derivative (1.0 eq.) and alkyne partner **3c**[1] **(**1.0 eq.) were dissolved in degassed tetrahydrofuran (THF) and stirred until completely dissolved. Degassed distilled water was added to the mixture, which was then stirred vigorously for an additional 10 minutes. To this mixture, freshly prepared aqueous solutions of sodium ascorbate (1.0 M, 0.05 eq.) and copper (II) sulfate ($CuSO_4$ 1.0 M, 0.10 eq.) were introduced in three equal portions at 45-min intervals. The reaction mixture was stirred at ambient temperature for 16 hours. Upon completion, the mixture was diluted with water and extracted with dichloromethane (DCM). The combined organic layers were dried over anhydrous sodium sulfate ($Na_2SO_4$), filtered, and concentrated under reduced pressure. The crude residue was purified via normal-phase silica gel column chromatography, eluting with a gradient of methanol in dichloromethane (MeOH/DCM), to yield the desired triazole conjugate **3d**. The previously obtained crude tosylate intermediate (1.0 eq.) was dissolved in anhydrous *N*,*N*-dimethylformamide (DMF) in a round-bottomed flask equipped with a magnetic stir bar. Sodium azide ($NaN_3$, 10.0 eq.) was added in one portion to the solution. The resulting heterogeneous mixture was stirred continuously at ambient temperature overnight. Upon completion, the reaction mixture was quenched by the careful addition of deionized water and transferred to a separatory funnel. The aqueous phase was thoroughly extracted with dichloromethane thrice. The organic layers were combined, washed with water and brine to remove residual DMF, and dried over anhydrous sodium sulfate $Na_2SO_4$. The drying agent was removed via filtration, and the filtrate was concentrated *in vacuo* under reduced pressure. The resulting crude material was purified by normal phase column chromatography on silica gel, utilizing a gradient elution system of ethyl acetate in hexanes (MeOH/DCM), to afford the pure azide **3e** target compound. To a stirred solution of the previously obtained compound (1.0 eq.) in anhydrous dichloromethane (DCM) was added trifluoroacetic acid (TFA, 4.0 eq.) dropwise. The reaction mixture was stirred at ambient temperature for 2 hours. Upon completion, the volatiles (excess TFA and DCM) were removed *in vacuo*. The resulting residue was diluted with ethyl acetate (EtOAc) and transferred to a separatory funnel. The organic layer was washed with brine, dried over anhydrous sodium sulfate ($Na_2SO_4$), filtered, and concentrated under reduced pressure. The crude residue was precipitated from diethyl ether ($Et_2O$) and filtered to afford compound **3f** as a white solid.

**Characterization of compounds:**

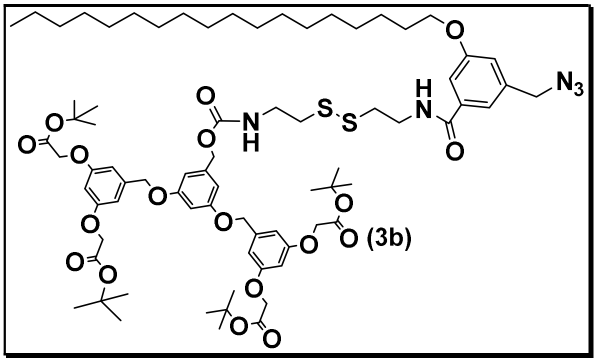

**Mol. formula:** $C_{76}H_{111}N_5O_{18}S_2$

**Mol. weight:** 1446.86 g/mol.

**Physical appearance:** White solid

**Yield:** 70%

To a stirred solution of compound **2h** (0.2g, 0.29mmol, 1.0 eq.) in anhydrous dichloromethane (DCM) taken in an oven-dried round-bottom flask, trifluoroacetic acid (TFA, 3.0 eq.) was added dropwise at room temperature. The reaction mixture was stirred at ambient temperature for 2 hours. Upon complete consumption of the starting material (monitored by TLC), the mixture was cooled to 0 °C and quenched via the careful addition of a saturated aqueous sodium bicarbonate ($NaHCO_3$) solution until the aqueous layer reached pH 8. The organic layer was separated, and the aqueous phase was extracted with DCM. The combined organic layers were dried over anhydrous sodium sulfate ($Na_2SO_4$), filtered, and concentrated under reduced pressure to afford the crude amine intermediate, which was utilized immediately in the subsequent step without further purification. The crude intermediate was dissolved in anhydrous DCM, and compound **1d** (0.26g, 0.25mmol, 0.8 eq.) was added to the solution. The mixture was stirred at room temperature for 10 minutes, followed by the dropwise addition of triethylamine ($Et_3N$, 0.1ml, 5.0 eq.). The resulting reaction mixture was allowed to stir for 4 hours. The mixture was then diluted with water, and the organic layer was separated. The organic phase was washed sequentially with saturated aqueous $NaHCO_3$ and brine, dried over anhydrous $Na_2SO_4$, filtered, and concentrated under reduced pressure. Purification of the crude residue via silica gel column chromatography using an ethyl acetate/hexane gradient (EtOAc/hexane) afforded the desired compound as white solid (0.26g, 0.18mmol) with 70% yield. $\mathbf{R_f}$ 0.45 in 35% EA/PE.

**$^{1}$H NMR (400 MHz, $CDCl_3$) δ** 7.30 (d, *J* = 19.6 Hz, 2H), 7.06 – 6.90 (m, 2H), 6.58 – 6.42 (m, 7H), 5.59 (t, *J* = 6.3 Hz, 1H), 4.96 (d, *J* = 40.4 Hz, 6H), 4.47 (s, 7H), 4.29 (s, 2H), 3.96 (t, *J* = 6.6 Hz, 2H), 3.77 – 3.69 (m, 2H), 3.52 (t, *J* = 6.4 Hz, 2H), 2.88 (d, *J* = 29.3 Hz, 4H), 1.75 (p, *J* = 6.7 Hz, 3H), 1.47 (s, 64H), 0.87 (t, *J* = 6.7 Hz, 4H).

**$^{13}$C NMR (100 MHz, $CDCl_3$) δ** 168.22, 167.62, 160.28, 160.10, 159.62, 139.88, 137.68, 118.85, 117.89, 113.36, 106.97, 106.90, 102.09, 101.71, 82.90, 70.16, 68.79, 66.77, 66.15, 54.78, 40.44, 39.48, 38.62, 38.28, 32.34, 30.11, 30.09, 30.07, 30.04, 30.01, 29.81, 29.77, 29.59, 28.46, 26.42, 23.10, 14.54.

**MALDI-TOF MS: M+K** 1485.23 g/mol.

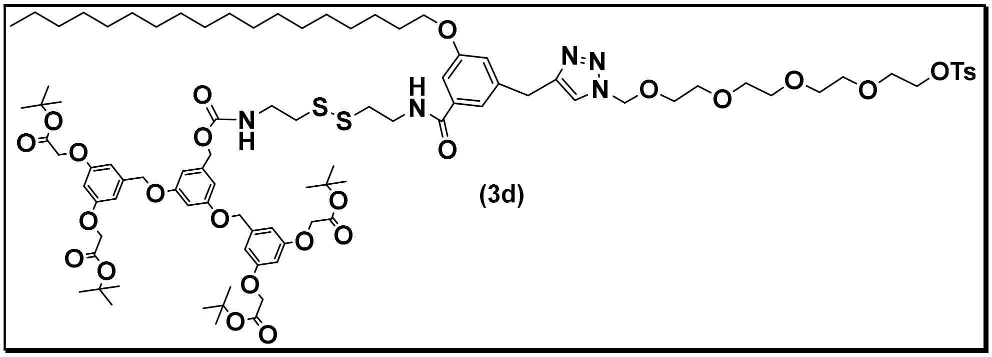


(3d)

**Mol. formula:** $C_{94}H_{137}N_5O_{25}S_3$

**Mol. weight:** 1833.32 g/mol

**Physical appearance:** Yellowish liquid

**Yield:** 85%

The azide derivative (80mg, 55µmol, 1.0 eq.) and alkyne **3c** (17.0mg, 44.23µmol, 0.8 eq.) were dissolved in degassed tetrahydrofuran (THF) and stirred until completely dissolved. Degassed distilled water was added to the mixture, which was then stirred vigorously for an additional 10 min. To this mixture, freshly prepared aqueous solutions of sodium ascorbate (1.0 M, 0.05 eq.) and copper (II) sulfate ($CuSO_4$ 1.0 M, 0.10 eq.) after 30 minutes. The reaction mixture was stirred at ambient temperature for 4 hours. Upon completion, the mixture was diluted with water and extracted with dichloromethane (DCM). The combined organic layers were dried over anhydrous sodium sulfate ($Na_2SO_4$), filtered, and concentrated under reduced pressure. The crude residue was purified via NPC, eluting with a gradient of methanol in dichloromethane (MeOH/DCM), to yield the desired product as yellowish liquid (86mg, 46.91 μmol) with 85% yield. **R$_f$** 0.5 in 5% MeOH/DCM

**$^1$H NMR (401 MHz, $CDCl_3$) δ** 7.76 (d, *J* = 7.9 Hz, 2H), 7.33 (d, *J* = 7.6 Hz, 2H), 6.89 (s, 1H), 6.62 – 6.35 (m, 8H), 4.90 (s, 4H), 4.47 (s, 8H), 4.11 (q, *J* = 7.1 Hz, 4H), 3.91 (s, 2H), 3.78 – 3.53 (m, 16H), 2.88 (d, *J* = 25.2 Hz, 4H), 2.43 (d, *J* = 6.7 Hz, 3H), 1.71 (d, *J* = 8.6 Hz, 3H), 1.47 (s, 35H), 1.28 – 1.23 (m, 36H), 0.88 (d, *J* = 6.3 Hz, 3H).

**MALDI TOF MS: M+Na** 1856.7 g/mol.

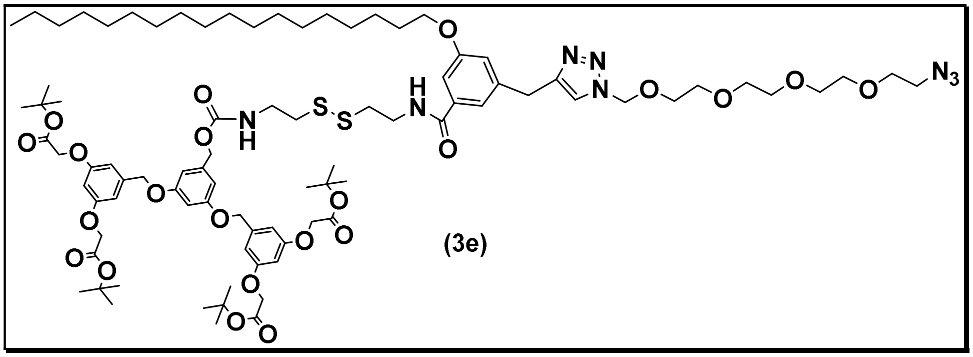

(3e)

**Mol. formula:** $C_{87}H_{130}N_8O_{22}S_2$

**Mol. weight:** 1704.15 g/mol.

**Physical appearance:** Yellowish liquid

**Yield:** 81%

The previously obtained tosylate intermediate 4b (75mg, 41μmol, 1.0 eq.) was dissolved in anhydrous *N*,*N*-dimethylformamide (DMF) in a round-bottomed flask equipped with a magnetic stir bar. Sodium azide ($NaN_3$, 26.60mg, 0.4mmol, 10.0 eq.) was added in one portion to the solution. The resulting heterogeneous mixture was stirred continuously at ambient temperature overnight. Upon completion, the reaction mixture was quenched by the careful addition of deionized water and transferred to a separatory funnel. The aqueous phase was thoroughly extracted with dichloromethane thrice. The organic layers were combined, washed with water and brine to remove residual DMF, and dried over anhydrous sodium sulfate $Na_2SO_4$. The drying agent was removed via filtration, and the filtrate was concentrated *in vacuo* under reduced pressure. The resulting crude material was purified NPC, utilizing a gradient elution system of methanol in dichloromethane MeOH/DCM, to afford the pure azide target compound as yellowish liquid (57mg, 33.45µmol) with 82% yield. **$R_f$** 0.5 in 5% MeOH/DCM

**$^1$H NMR (400 MHz, $CDCl_3$) δ** 7.35 (s, 1H), 7.07 – 6.87 (m, 1H), 6.65 – 6.34 (m, 8H), 4.95 (d, *J* = 38.5 Hz, 5H), 4.48 (s, 7H), 3.92 (s, 2H), 3.65 (d, *J* = 16.9 Hz, 15H), 3.36 (s, 2H), 2.90 (dd, *J* = 29.1, 9.7 Hz, 4H), 1.72 (d, *J* = 7.4 Hz, 2H), 1.47 (s, 35H), 1.26 (d, *J* = 12.5 Hz, 35H), 0.86 (d, *J* = 7.0 Hz, 4H).

**$^{13}$C NMR (101 MHz, $CDCl_3$) δ** 168.21, 160.27, 159.61, 139.86, 136.68, 118.21, 113.81, 106.93, 102.04, 101.68, 82.89, 71.09, 71.04, 71.00, 70.96, 70.44, 70.33, 70.18, 68.90, 66.15, 51.13, 32.33, 32.05, 30.72, 30.11, 30.06, 30.01, 29.92, 29.82, 29.77, 29.57, 28.47, 26.41, 23.10, 14.54.

**MALDI TOF MS: M+Na** 1726.5 g/mol.

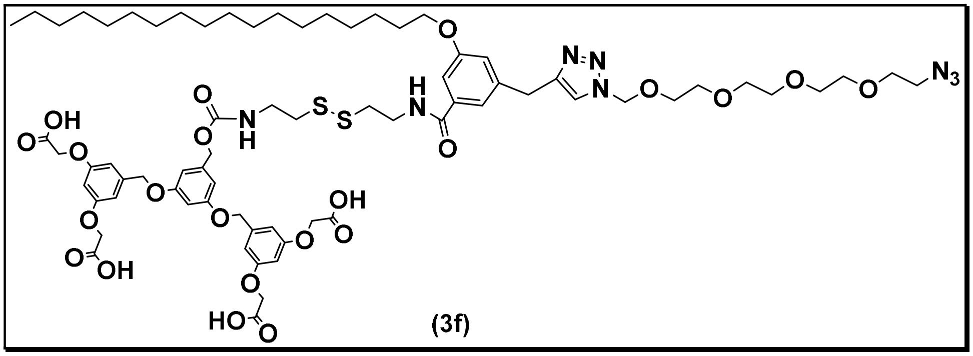


(3f)

**Mol. formula:** $C_{71}H_{98}N_8O_{22}S_2$

**Mol. weight:** 1478.62 g/mol.

**Physical appearance:** White solid

**Yield:** 77%

To a stirred solution of the previously obtained intermediate **3c** (1.0 eq.) in anhydrous dichloromethane (DCM) was added trifluoroacetic acid (TFA, 4.0 eq.) dropwise at room temperature. The reaction mixture was stirred continuously at ambient temperature for 2 hours, during which complete consumption of the starting material was confirmed by TLC analysis. After completion of the reaction, the solvent and excess TFA were removed under reduced pressure using a rotary evaporator to afford a crude residue. The residue was dissolved in ethyl acetate (EtOAc) and transferred to a separatory funnel. The organic phase was washed with saturated aqueous sodium chloride (brine) to remove residual acid and water-soluble impurities, then dried over anhydrous sodium sulfate ($Na_2SO_4$). The drying agent was removed by filtration, and the filtrate was concentrated under reduced pressure to obtain the crude product. The resulting residue was triturated with cold diethyl ether ($Et_2O$), leading to the formation of a white precipitate. The solid was collected by vacuum filtration, washed with additional cold $Et_2O$, and dried under high vacuum to afford compound **3d** as a white solid.

**$^1$H NMR (401 MHz, DMSO-$d_6$) δ** 8.65 (s, 1H), 8.13 (s, 1H), 7.50 – 7.27 (m, 3H), 7.01 (s, 1H), 6.60 (d, $J$ = 2.2 Hz, 6H), 6.42 (d, $J$ = 2.3 Hz, 2H), 5.56 (s, 2H), 4.96 (d, $J$ = 13.0 Hz, 6H), 4.65 (s, 8H), 4.50 (s, 3H), 2.90 – 2.85 (m, 2H), 2.81 (d, $J$ = 6.8 Hz, 2H), 1.67 (d, $J$ = 7.2 Hz, 2H), 1.22 (s, 30H), 0.83 (s, 3H).

**$^{13}$C NMR (101 MHz, DMSO-$d_6$) δ** 170.35, 166.01, 159.76, 159.21, 159.11, 156.36, 144.67, 139.80, 139.63, 137.89, 124.38, 119.49, 117.41, 112.72, 106.75, 100.88, 70.11, 70.08, 70.05, 69.98, 69.54, 69.38, 69.32, 68.10, 64.90, 63.79, 60.11, 52.84, 50.29, 46.07, 31.60, 29.32, 29.30, 29.07, 29.01, 28.89, 25.78, 22.40, 21.07, 14.38, 14.25, 8.90.

**MALDI TOF MS: M+Na** 1502.2 g/mol.

#### 4. NMR Data:

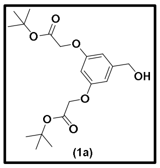

(1a)

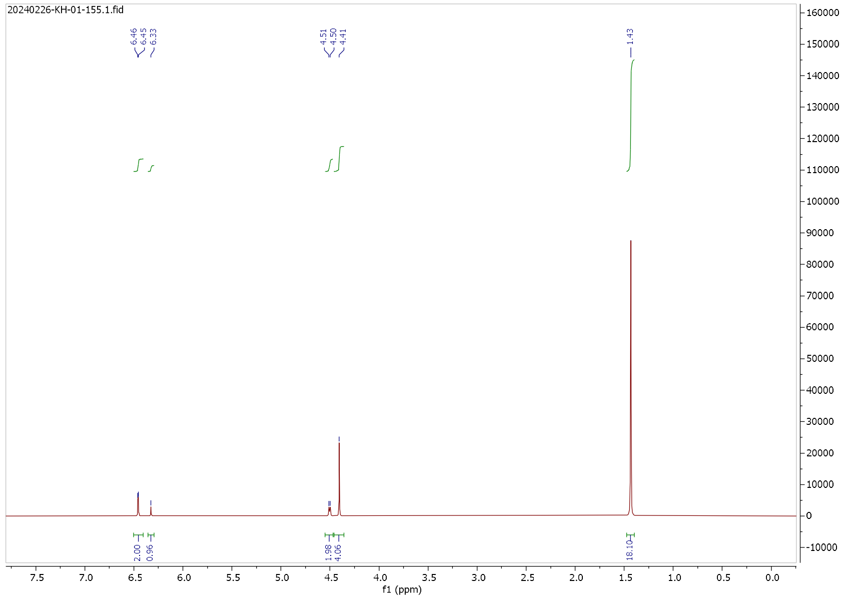


**$^{1}$H NMR Spectrum**

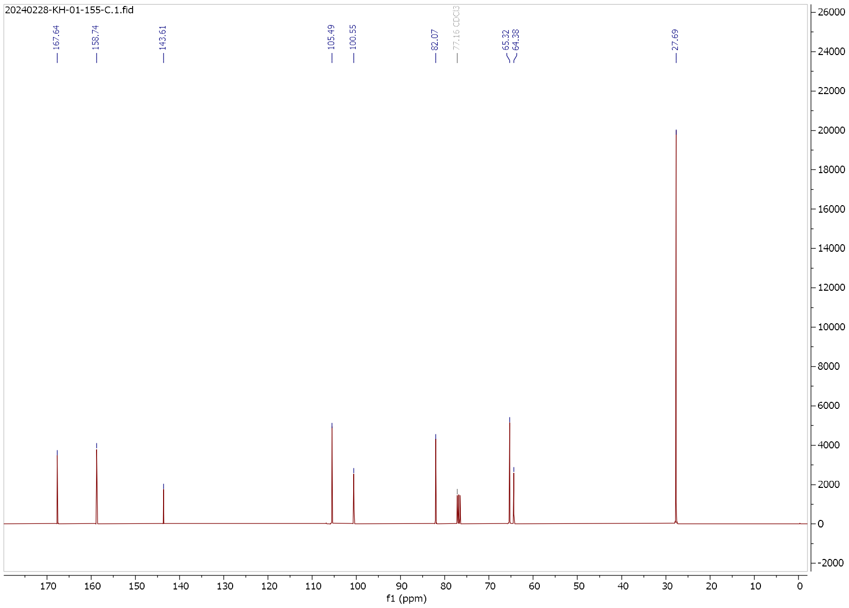


**$^{13}$C NMR Spectrum**

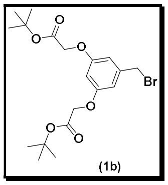


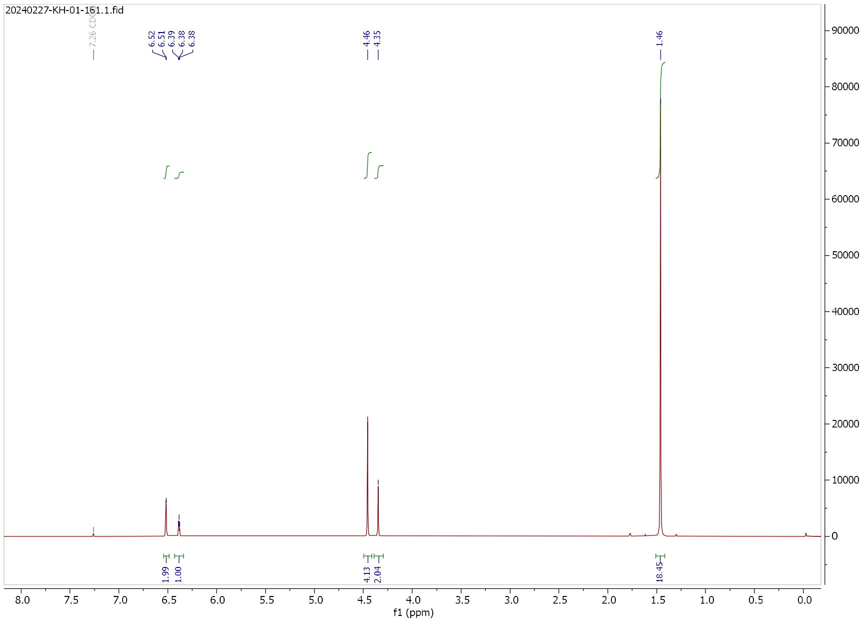

### $^{1}$H NMR Spectrum

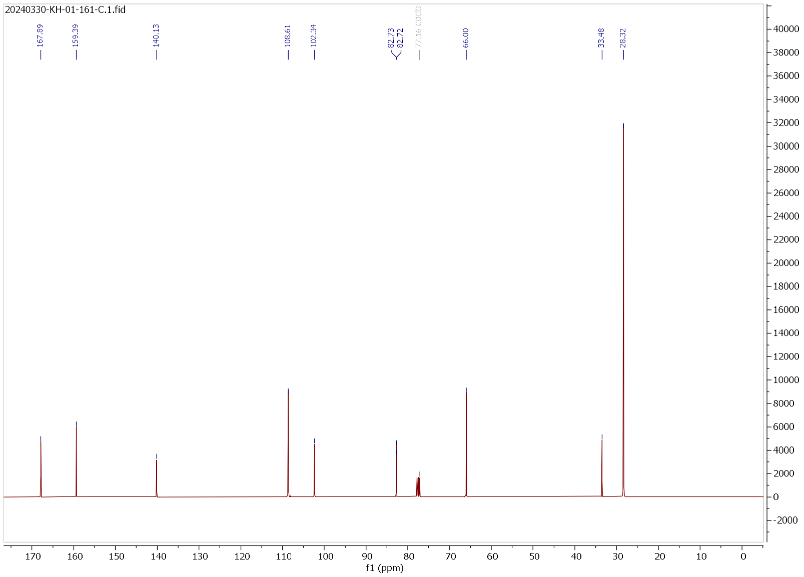

### $^{13}$C NMR Spectrum

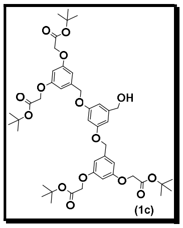


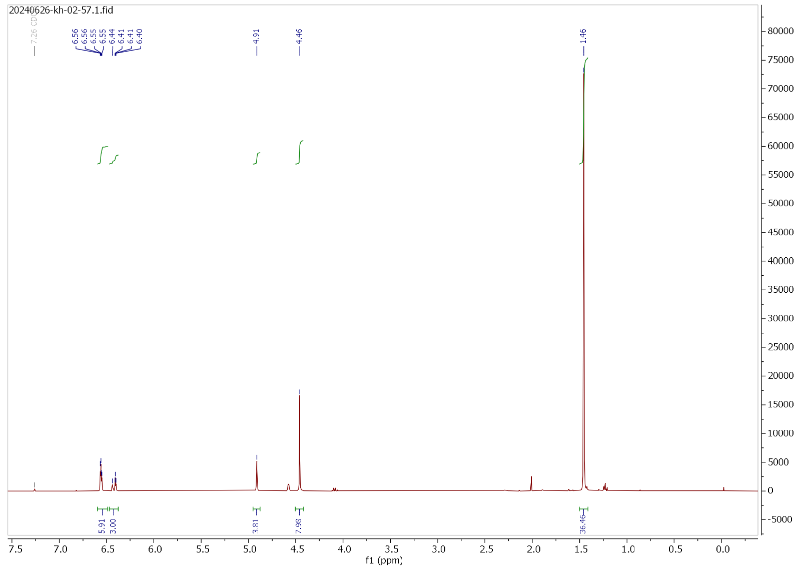

### $^{1}$H NMR Spectrum

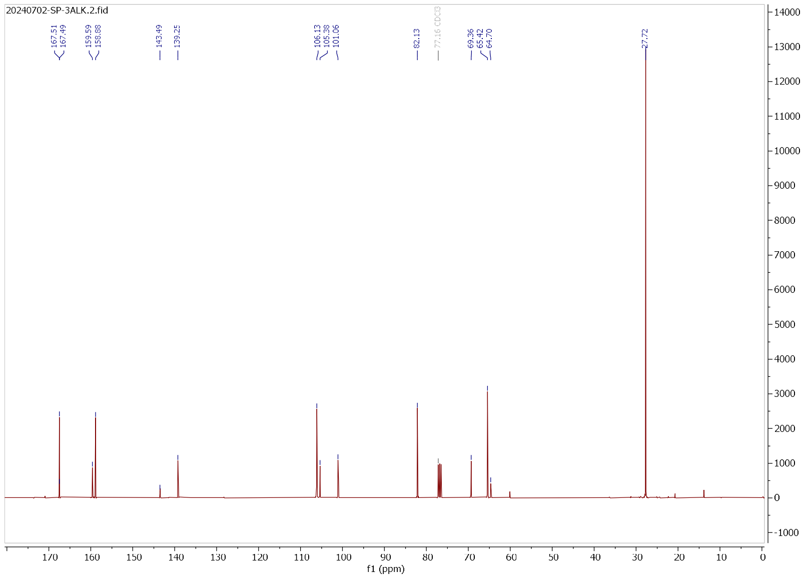

### $^{13}$C NMR Spectrum

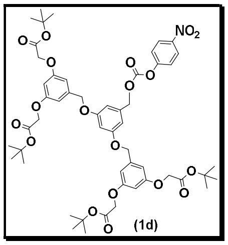


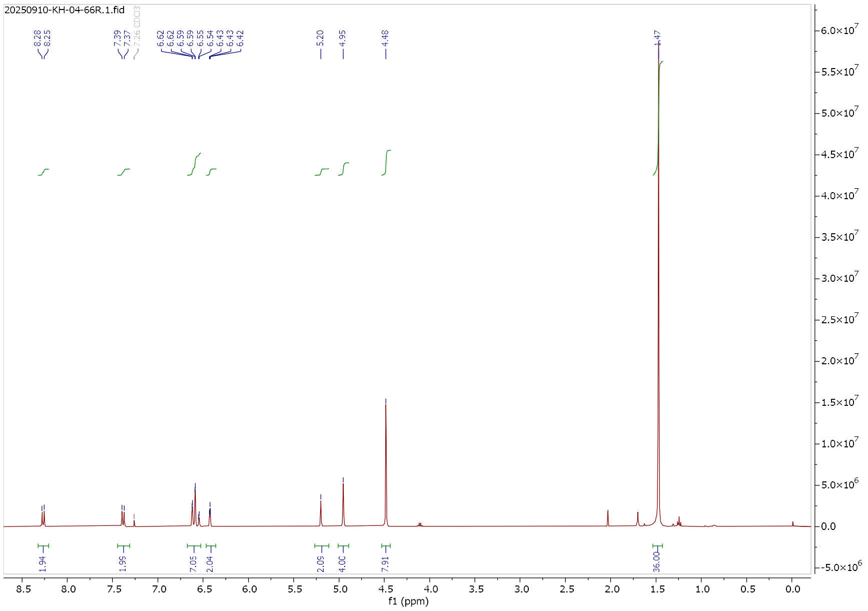


#### [1]H NMR Spectrum

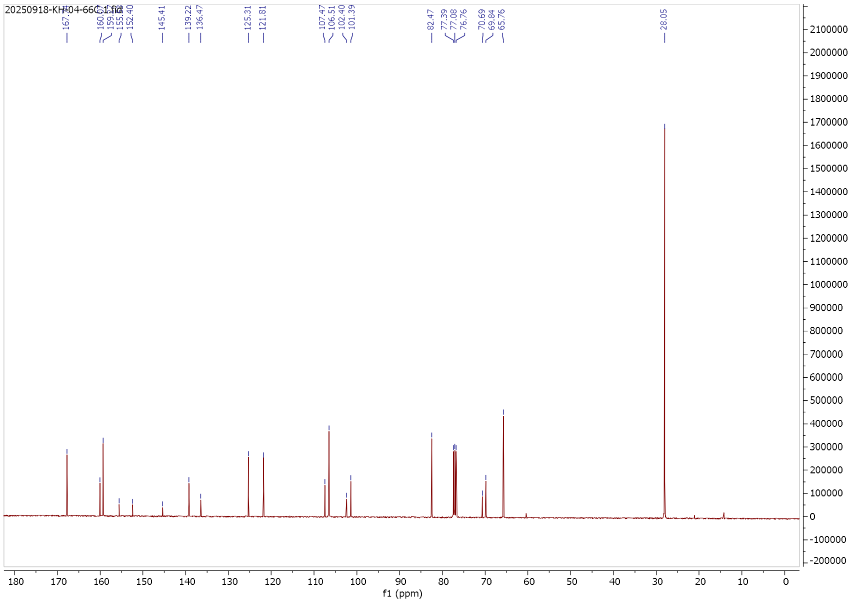


#### [13]C NMR Spectrum

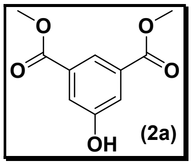


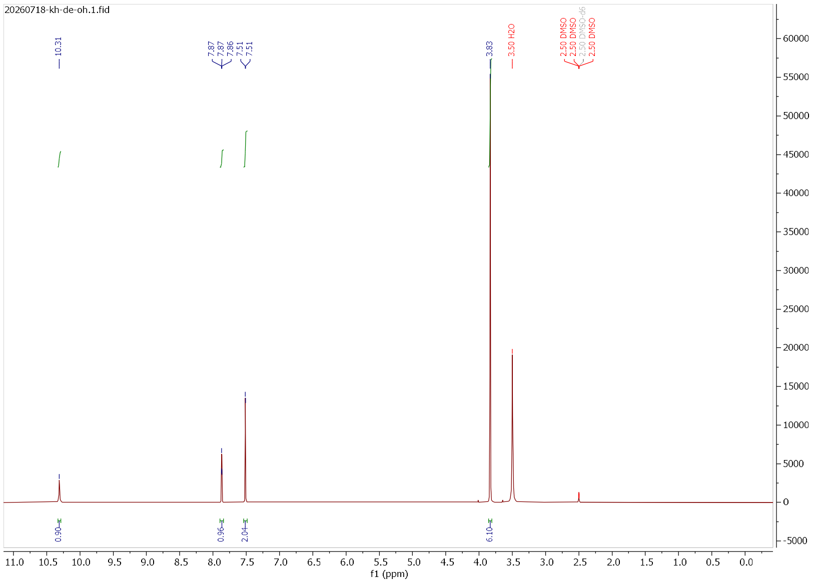


#### $^{1}H$ NMR Spectrum

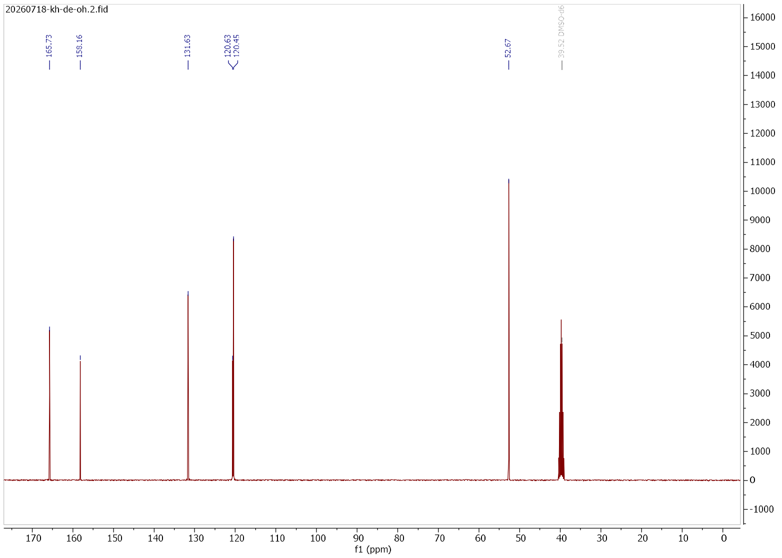


#### $^{13}C$ NMR Spectrum

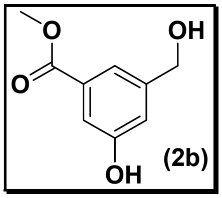


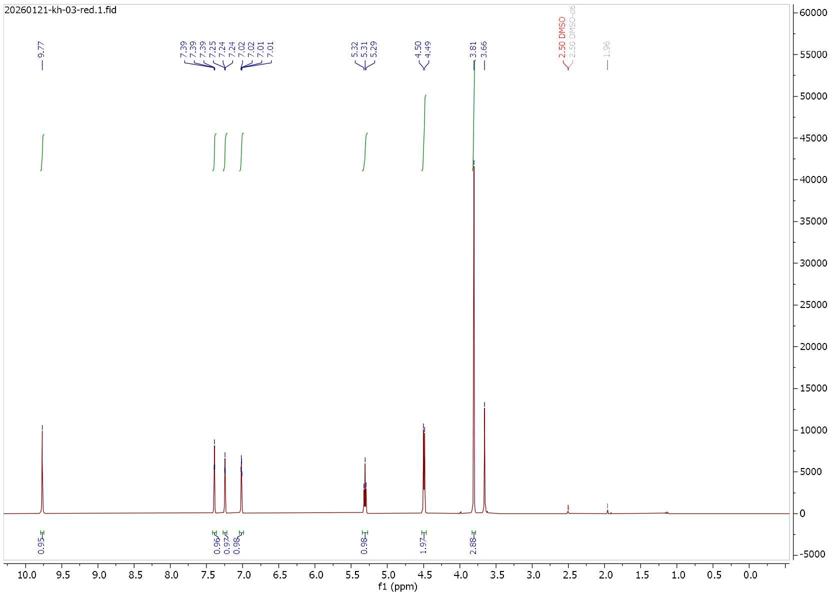


#### $^{1}$H NMR Spectrum

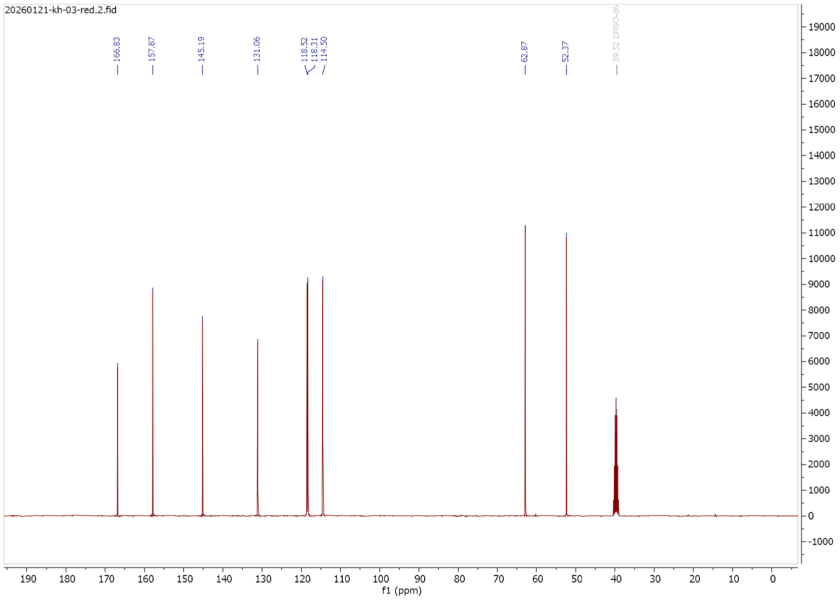


#### $^{13}$C NMR Spectrum

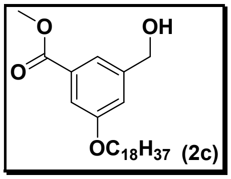

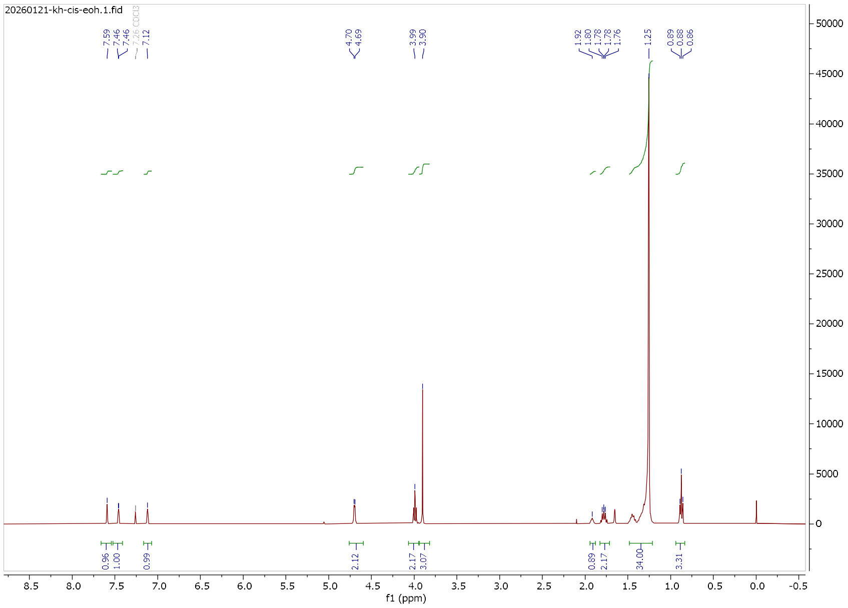

**$^{1}$H NMR Spectrum**

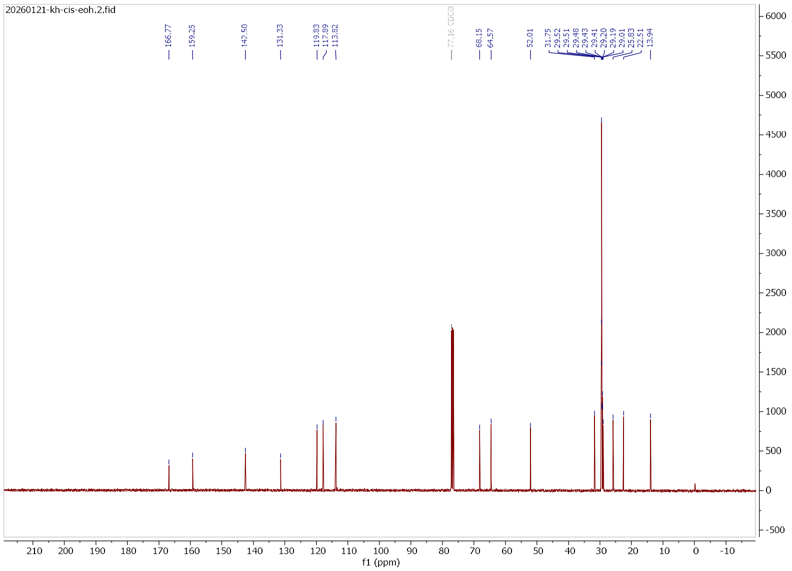

**$^{13}$C NMR Spectrum**

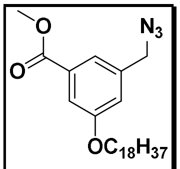


(2e)

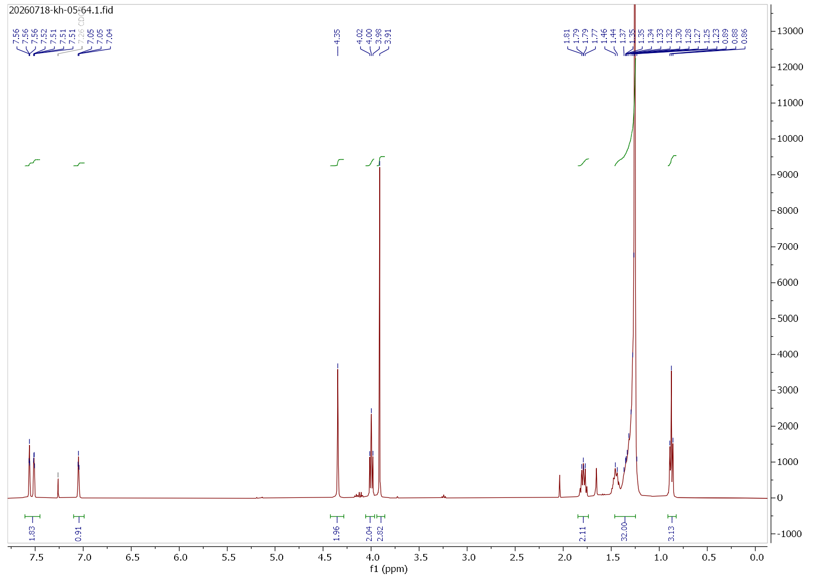


**$^1$H NMR Spectrum**

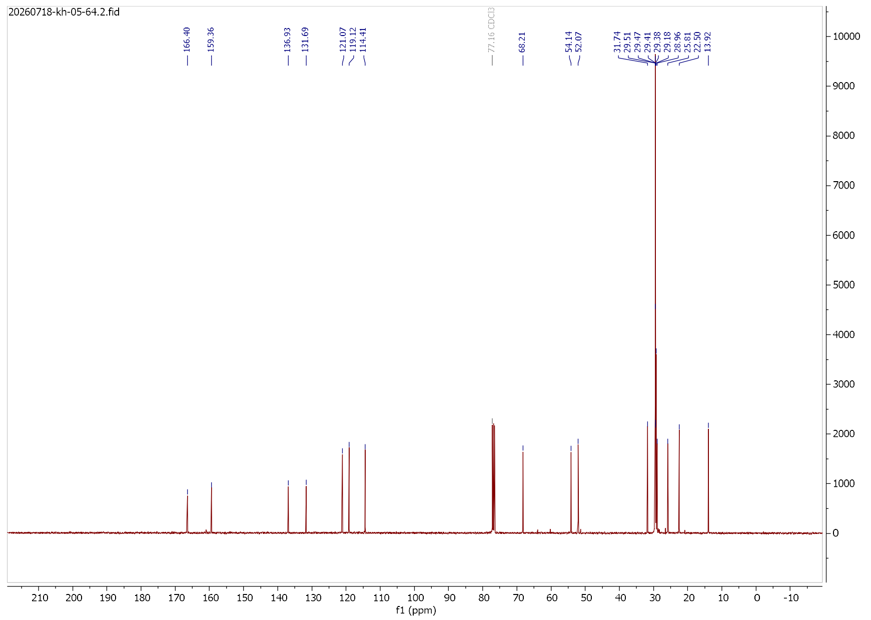


**$^{13}$C NMR Spectrum**

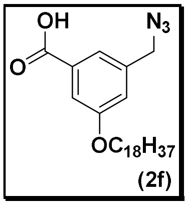


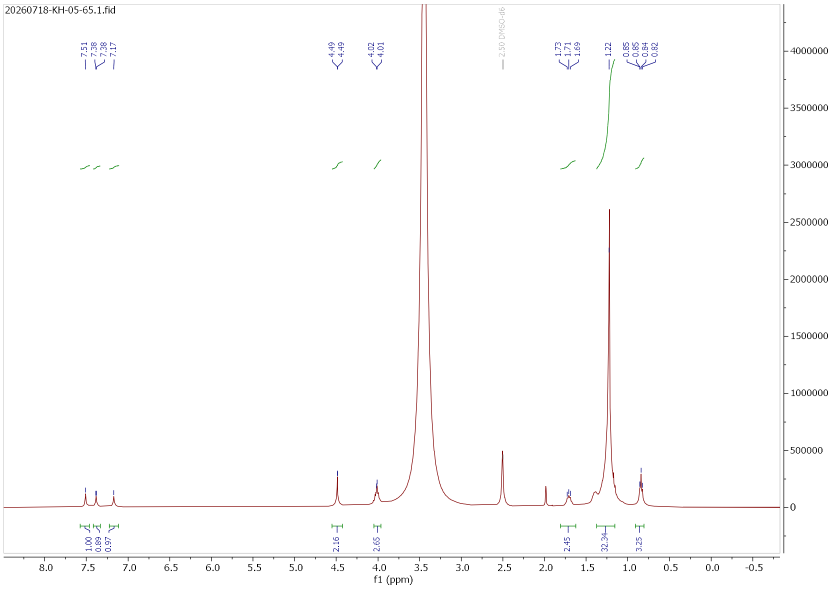


### $^{1}H$ NMR Spectrum

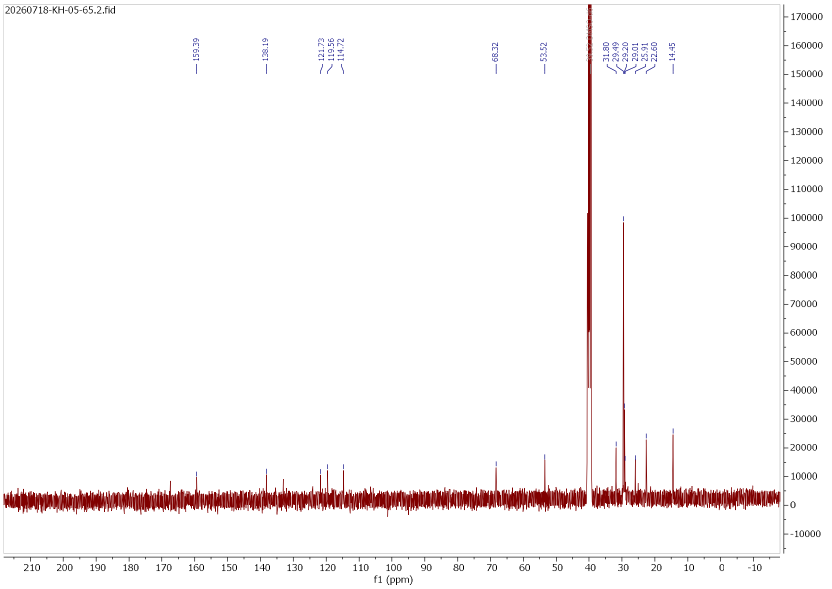


### $^{13}C$ NMR Spectrum

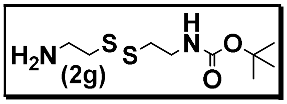

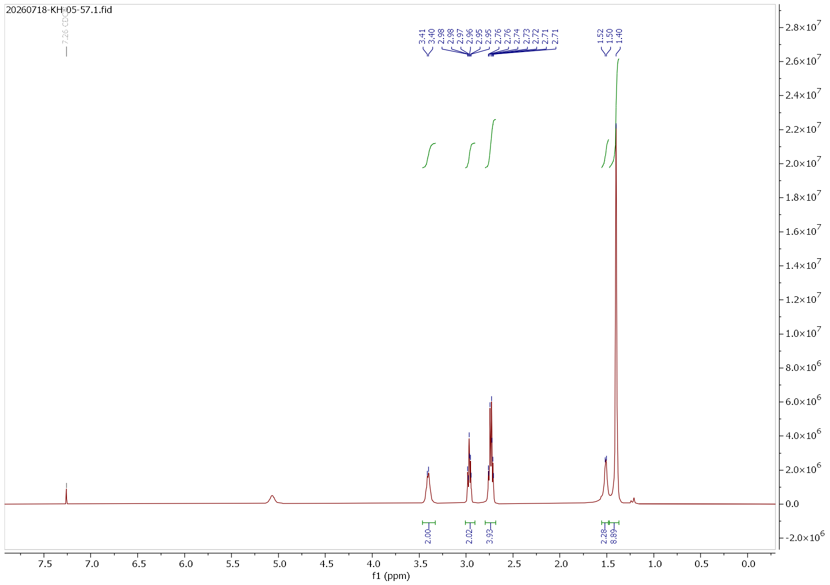

**$^{1}$H NMR Spectrum**

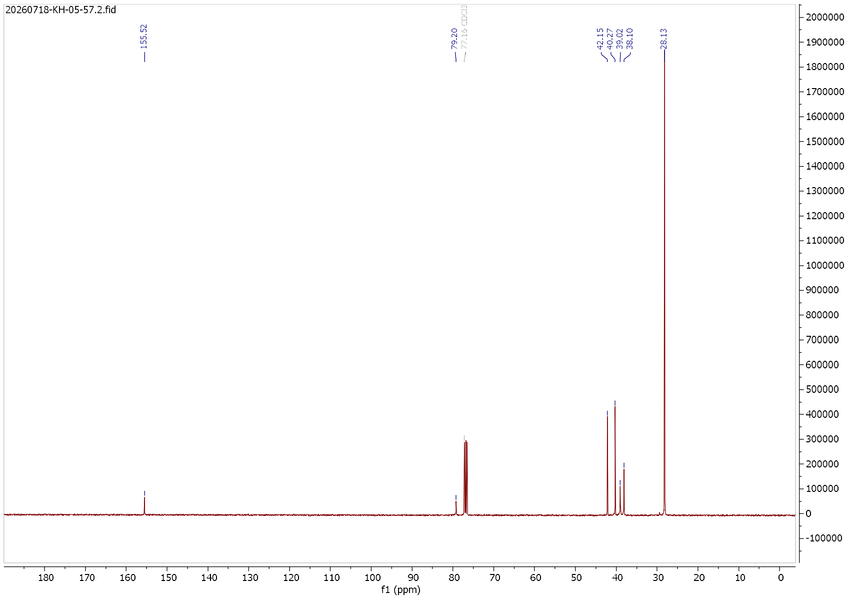

**$^{13}$C NMR Spectrum**

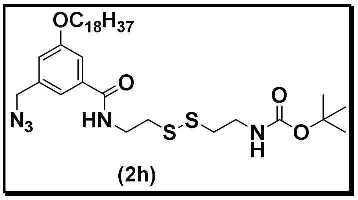

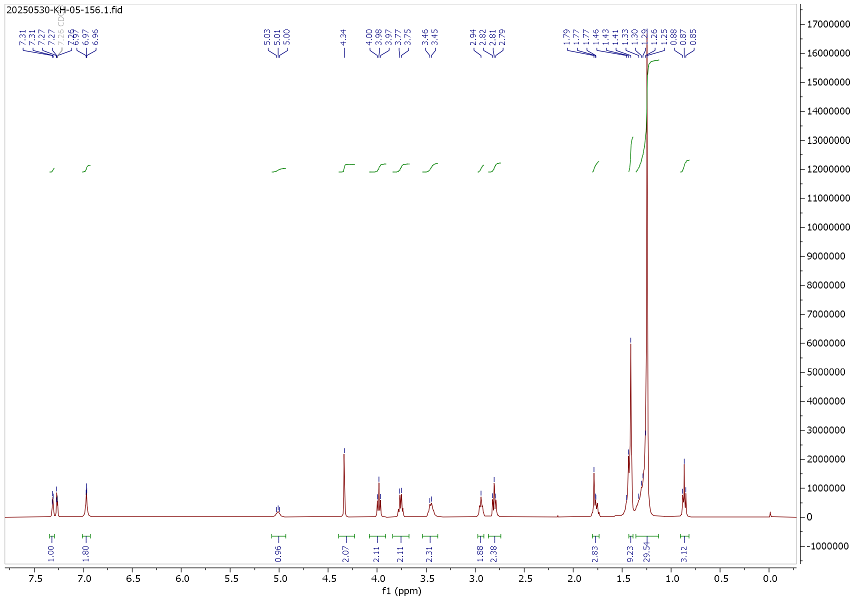

### $^{1}H$ NMR Spectrum

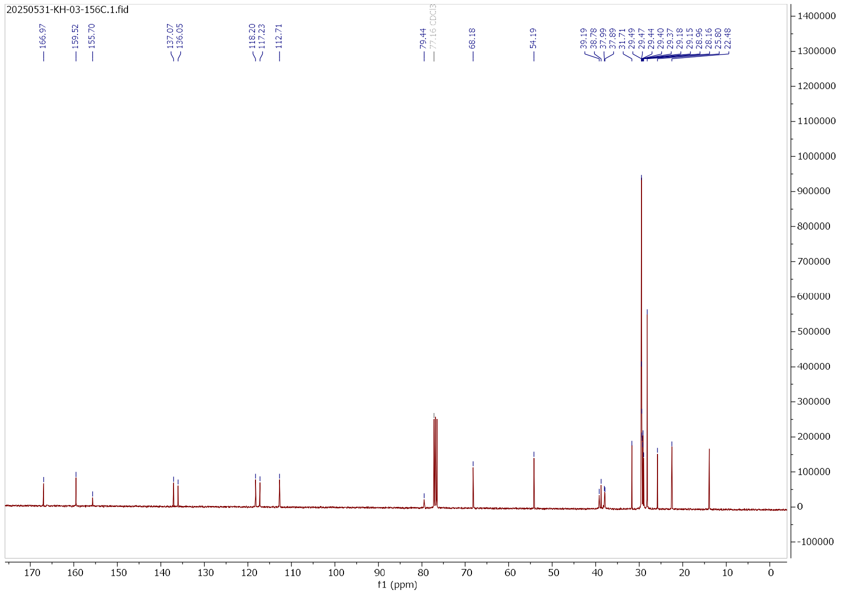

### $^{13}C$ NMR Spectrum

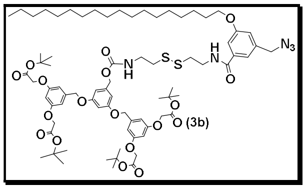


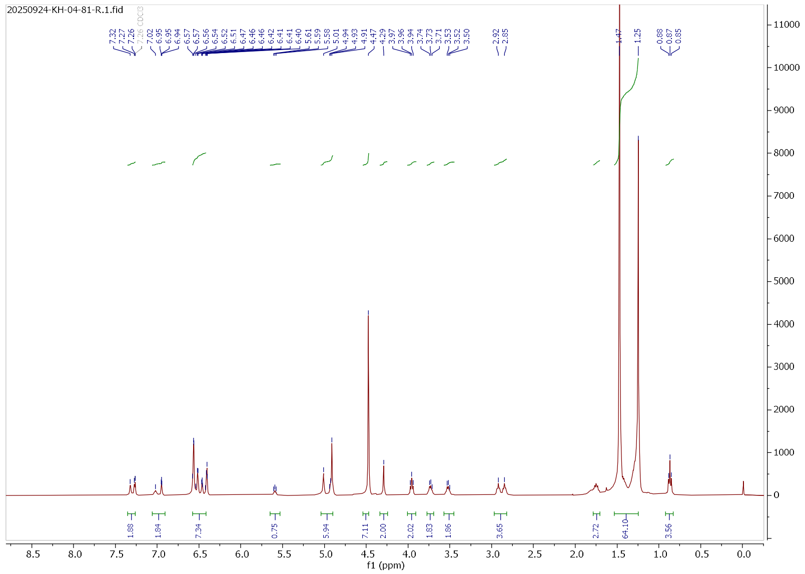


**¹H NMR Spectrum**

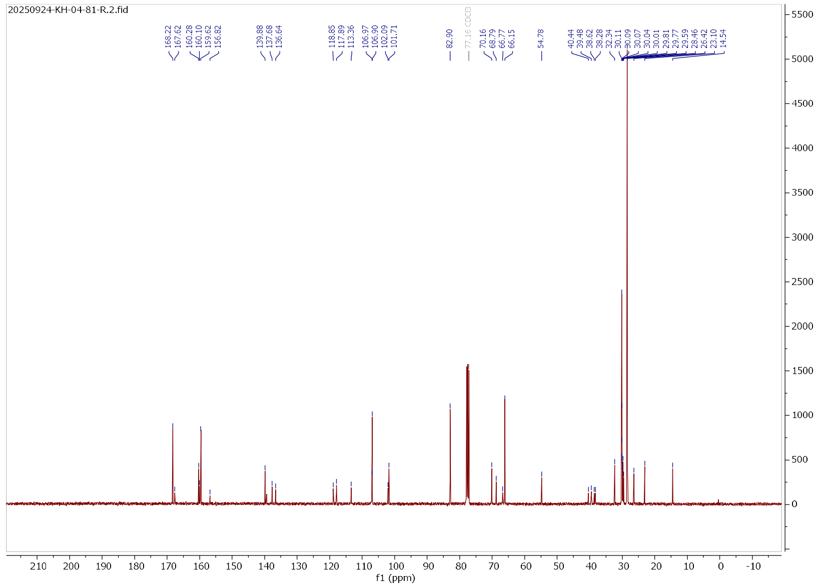


**¹³C NMR Spectrum**

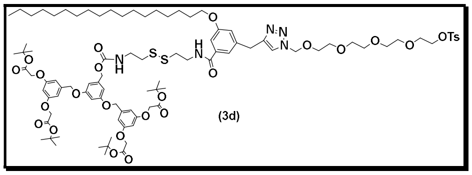

(3d)

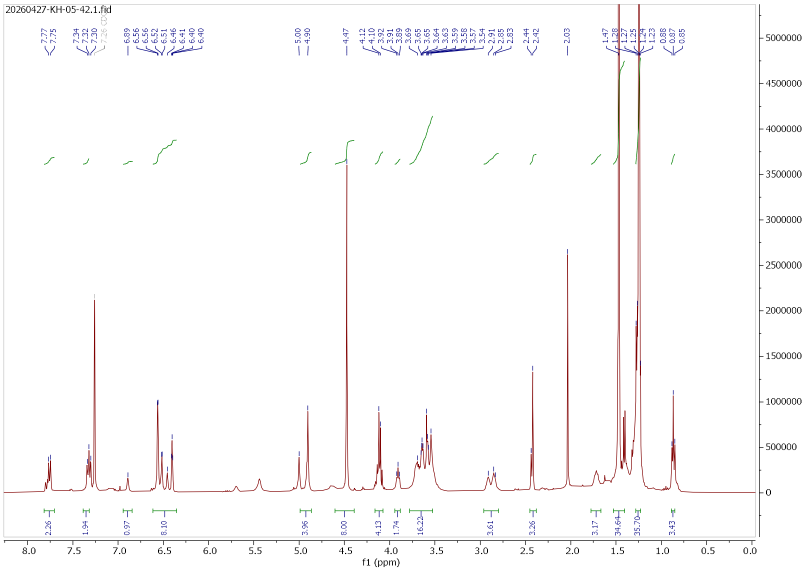


**$^{1}$H NMR Spectrum**

Spectrum Report

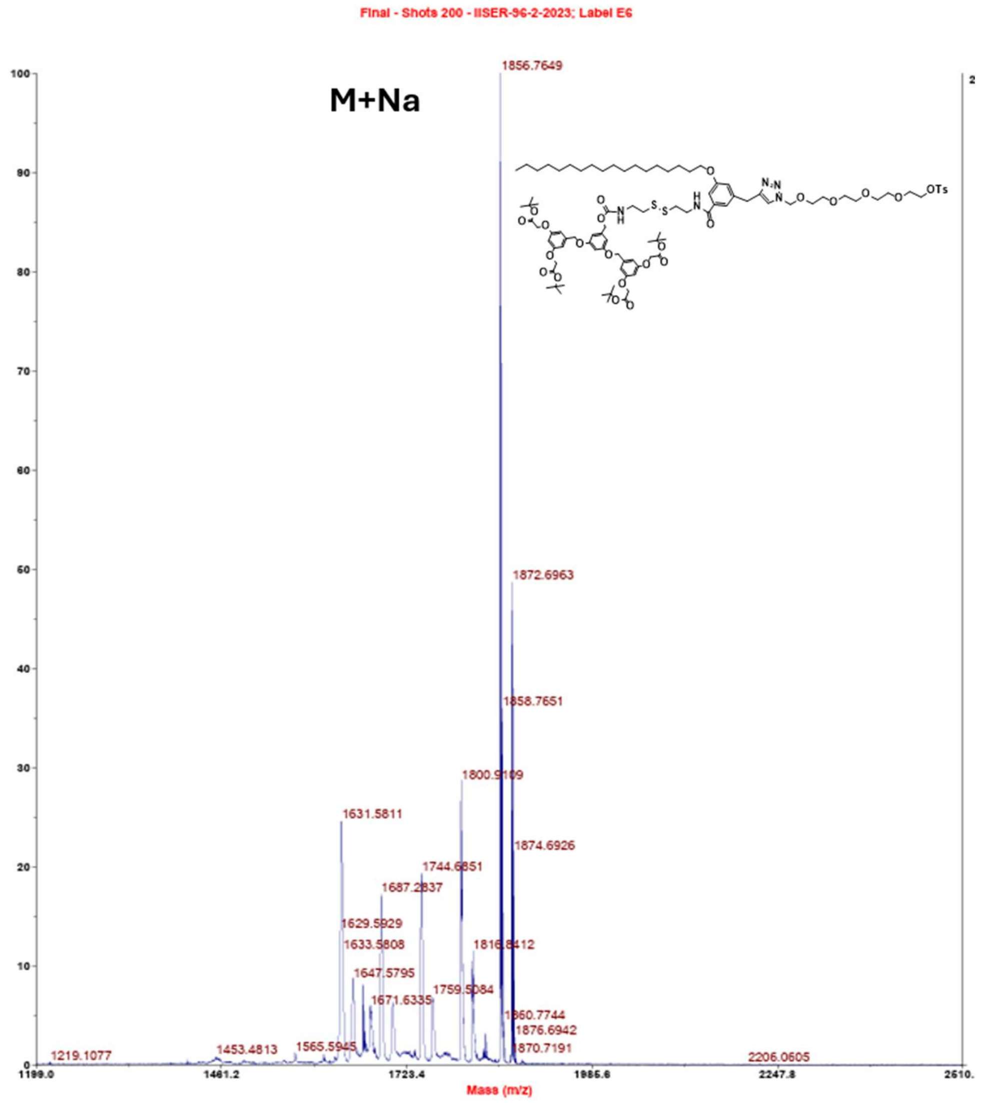


:ER-2023\IISER-96-2-2023 Label E6 Run # 669 Page 1 4/27/2026 5:02:5

**MALDI TOF M/S for compound 3d**

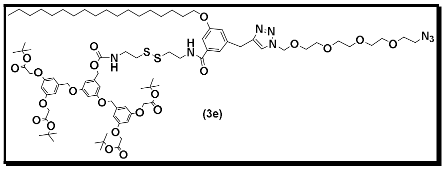

(3e)

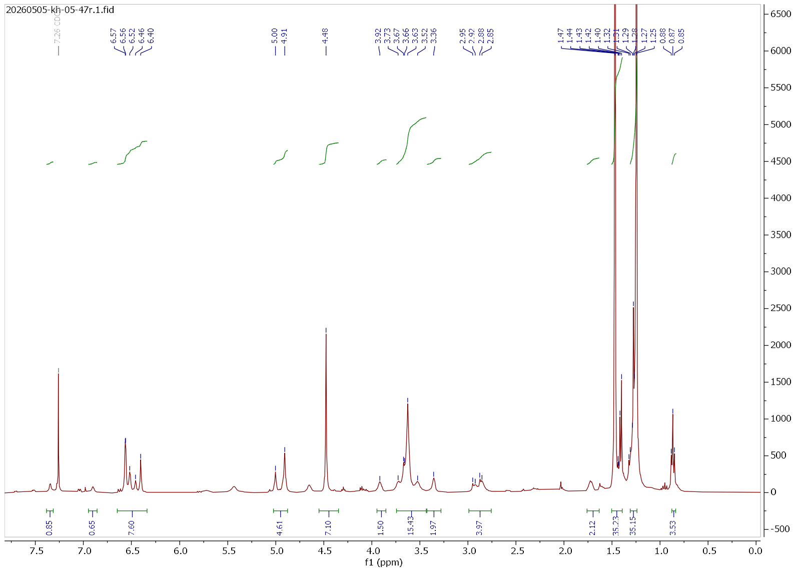


### [1]H NMR Spectrum

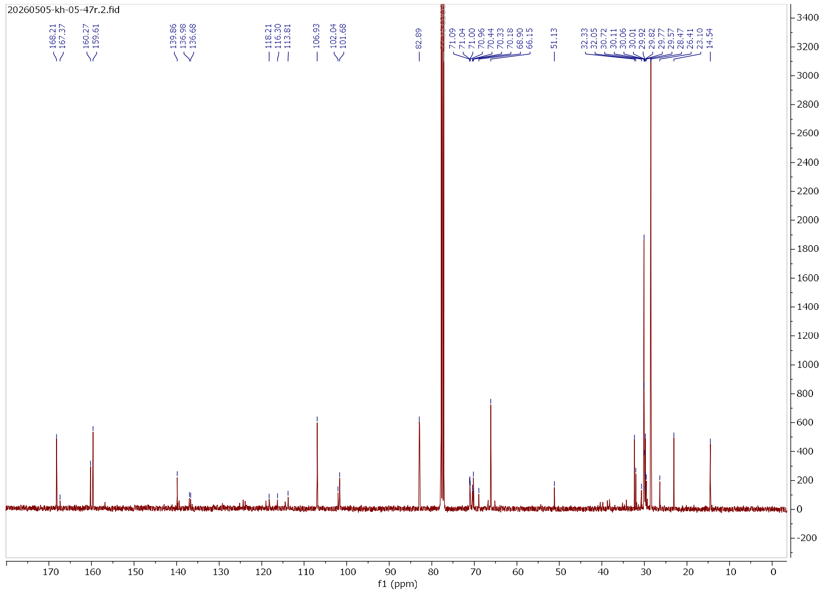


### [13]C NMR Spectrum

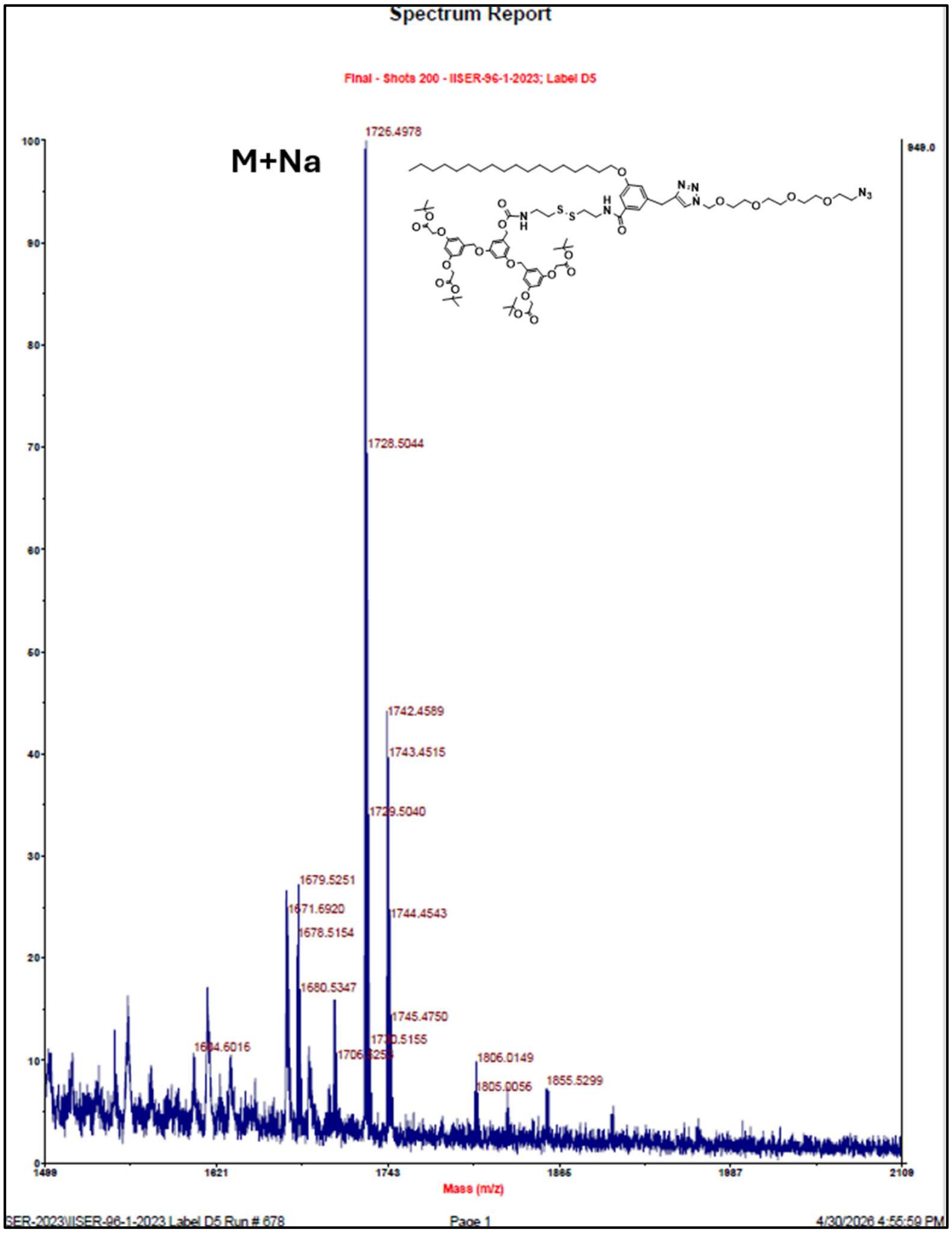


**MALDI TOF M/S for compound 3e**

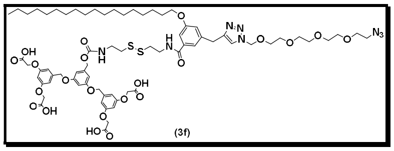


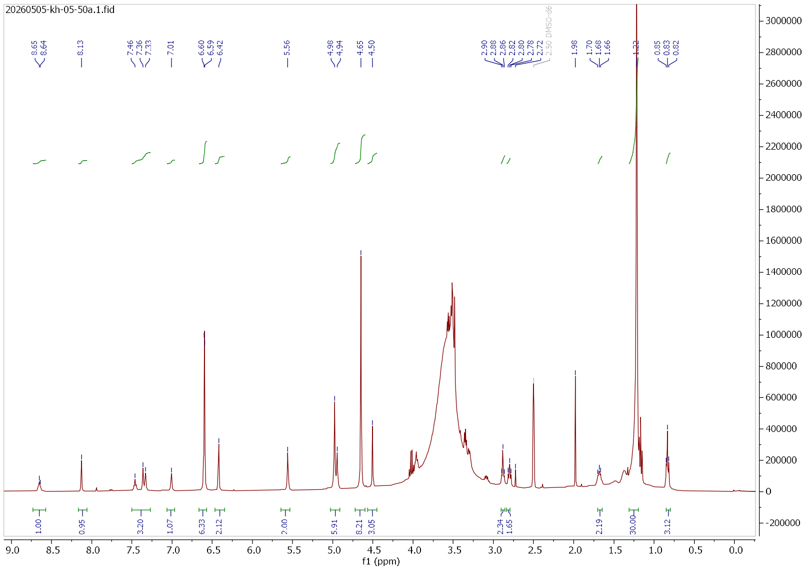

### $^{1}H$ NMR Spectrum

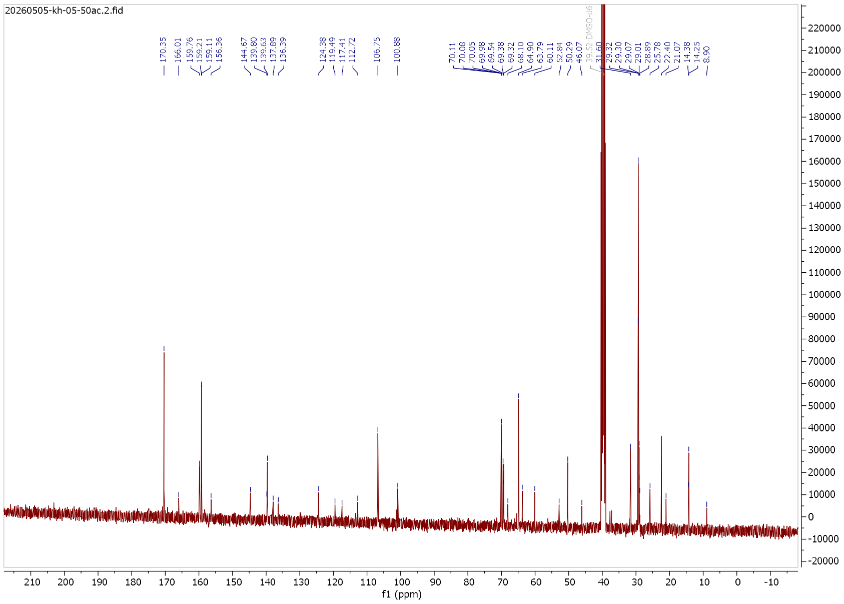

### $^{13}C$ NMR Spectrum

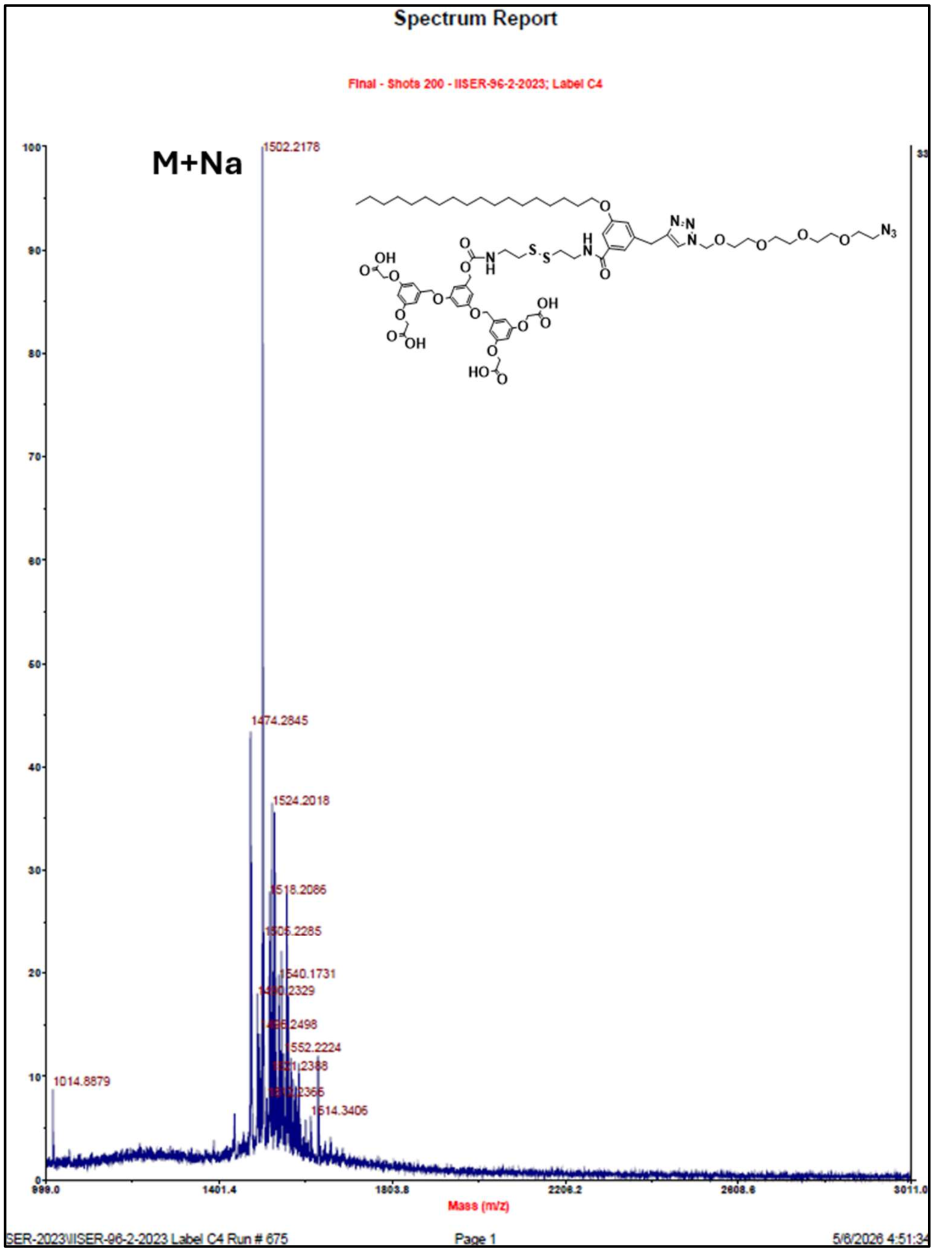


MALDI TOF M/S for compound 3f
